# When Does Active Learning Improve ADMET Prediction? A Multi-Endpoint Benchmark

**DOI:** 10.64898/2026.10.26.761649

**Authors:** Viktor Najdovski, Jana Hristova, Viktorija Vodilovska, Ilinka Ivanoska

## Abstract

Experimental ADMET profiling is essential to drug discovery but incredibly costly that only a fraction of candidate molecules can be measured, motivating active learning to prioritize informative compounds. However, evidence remains limited on how acquisition strategies and uncertainty estimators interact across different ADMET tasks. Thus, we present a protocol-matched benchmark of seven acquisition strategies and two uncertainty approaches (Deep Ensembles and MC Dropout) across eight ADMET regression endpoints. Our framework combines molecular prediction, uncertainty estimation, batch aquisition and empirically calibrated prediction intervals, with a strict chemotype split to evaluate generalization to held-out chemical groups. The results reveal that active-learning benefits were endpoint- and strategy-dependent, with the broadest favorable pattern observed for solubility and diversity-aware approaches showing the most consistently favorable behavior across endpoints. Deep Ensembles achieved lower mean RMSE than MC Dropout in 45 of 56 matched comparisons, although this predictive advantage did not consistently translate into better acquisition. Post-hoc analyses associated this disconnect with weak uncertainty-error alignment and reduced chemotype coverage in selected batches. This benchmark offers a reproducible basis for choosing endpoint-specific acquisition strategies, assessing prediction quality alongside computational cost, and designing prospective studies with matched random controls.

## Introduction

Before a drug candidate can enter clinical trials, it must pass a battery of laboratory tests to determine its ADMET (absorption, distribution, metabolism, excretion, toxicity) profile ^1^. Failures at this stage are both common and costly. Unfavorable pharmacokinetics and toxicity have historically accounted for a substantial share of clinical attrition and remain among the dominant non-efficacy reasons that candidates are abandoned after heavy investment^2,3^. Moving that characterization earlier is the standard response, and it is effective, but it is not cheap. A typical pharmaceutical library contains millions of candidates, while a single ADMET assay, requiring a purified compound, specialized cell lines or microsomes, and liquid-handling equipment, can cost tens to hundreds of dollars per molecule and take days to weeks per batch ^4^. Experimental budgets therefore permit only a small fraction of a library to be measured, which makes the choice of that fraction a design decision in its own right rather than an afterthought.

Predictive models are already load-bearing infrastructure in this workflow. Large pharmaceutical organizations maintain in-house ADMET platforms that serve predictions continuously and are retrained as new measurements accumulate ^5^. Their sustained value depends on training data that keeps pace with the chemistry under exploration. A prospective industrial evaluation spanning 120 internal datasets and six *in vitro* endpoints found that model rankings established retrospectively do not always survive contact with genuinely new compounds, and that relative performance drifts as project chemistry moves ^6^. Recent ADMET benchmarking points in the same direction, reporting that data representation and evaluation design matter more than architectural sophistication, with standard cheminformatics representations remaining competitive against learned ones ^7^. A major practical constraint in this domain is which molecules carry reliable labels, in addition to model expressiveness ^8^.

Machine learning offers a way to extend a limited set of measurements to unmeasured compounds ^9^. The field has moved from hand-crafted descriptors paired with classical regressors such as random forests and support-vector machines ^10^ and from circular fingerprints ^11^ to message-passing networks that learn task-specific representations directly from the molecular graph ^12,13^. Reported accuracies on public benchmarks are frequently high, but they are typically obtained under random splits that place structurally similar analogues on both sides of the partition. Because medicinal chemistry series are assembled around a comparatively small number of recurring molecular frameworks ^14^, such splits reward interpolation within a scaffold rather than transfer between scaffolds, and scaffold- or time-based evaluations consistently show that random-split estimates overstate prospective performance ^15^. Reliable generalization to genuinely new chemotypes still requires large, chemically diverse training sets, which is precisely what limited assay throughput denies. Active learning (AL) addresses the constraint directly by choosing which molecules to measure ^16^. In the pool-based batch setting used here, a model is trained on a small initial labeled set, scores an unlabeled pool, acquires the most informative batch for measurement, and retrains on the enlarged set. If the selection criterion is well calibrated, the model should reach a target accuracy with far fewer labels than random sampling would require. The idea has been applied across molecular discovery, from accelerating virtual screening over very large libraries ^17^ to guiding iterative experimental design ^18^, and it is attractive in ADMET precisely because each avoided assay is a direct saving in reagents, instrument time, and compound.

That expectation deserves closer scrutiny than it usually receives, because active learning’s empirical record outside chemistry is mixed. Controlled re-evaluations in natural language processing found that the advantage of acquisition over random sampling does not transfer reliably across models and tasks, and can even invert when the underlying model is changed ^19^. A systematic study in computer vision reached a sharper conclusion that once sources of stochasticity are controlled and regularization is held fixed, a range of uncertainty-based, diversity-based, and committee-based acquisition functions offer marginal or no advantage over a random baseline, and several previously published gains did not reproduce ^20^. Both converge on the same methodological point that random acquisition under a matched budget is a stronger baseline, and a more consequential one to omit, than the literature generally assumes. The closest evidence from chemistry sharpens the point that active-learning benefits depend on both the chemical distribution and acquisition strategy. In a systematic study of aqueous solubility and redox potential, uncertainty-guided acquisition did not outperform random sampling for in-distribution compounds. Improvements were limited to structurally novel regions and were largest for a density-based selection criterion ^21^. *These findings motivate evaluating whether acquisition strategies remain beneficial across multiple ADMET endpoints under structural distribution shift*.

There are concrete reasons to expect ADMET to be a demanding case. Pure uncertainty sampling tends to select structurally redundant compounds from the same sparse region of chemical space, spending an entire assay plate on near-duplicates. Wet-lab measurements also carry irreducible experimental noise (aleatoric uncertainty). If an acquisition method cannot distinguish this aleatoric uncertainty from epistemic uncertainty caused by insufficient training data, it may repeatedly select compounds with inherently noisy measurements rather than compounds that expand the model’s knowledge of chemical space ^22^. ADMET endpoints further differ sharply in their marginal distributions, spanning near-symmetric physicochemical properties and clearance values covering several orders of magnitude, so a single modeling and evaluation strategy does not transfer cleanly across tasks. These considerations motivate assessing uncertainty quality together with the chemical diversity of selected batches.

Uncertainty-based active learning also poses methodological challenges. Uncertainty quality is itself an open research question in drug design ^23^. Information-theoretic acquisition based on the mutual information between predictions and model parameters ^24^ was extended to deep networks through MC dropout ^25^, and deep ensembles have since become the competing standard. Systematic comparisons for molecular property prediction find that calibration quality varies considerably across estimator and endpoint^26^, and conformal prediction has been advocated in cheminformatics because it can supply finite-sample coverage guarantees under exchangeability ^27^, including in combination with test-time dropout for deep networks ^28^. However, standard conformal coverage guarantees assume that calibration and test compounds are exchangeable^29^.

Against this background, we investigate whether active-learning strategies provide reproducible benefits over random acquisition across multiple different ADMET endpoints and under structural distribution shift. Specifically, we address the following research questions:

1. How does the performance of active-learning strategies relative to matched random acquisition vary across ADMET endpoints?
2. How do MC Dropout and Deep Ensembles differ in predictive performance and acquisition effectiveness?
3. Are acquisition outcomes associated with uncertainty-error alignment and the chemotype coverage of the selected batches?

To answer these questions, we benchmark six active-learning strategies and a matched random baseline across eight OpenADMET ExpansionRx ^30^ regression endpoints. Epistemic uncertainty is estimated using MC Dropout or Deep Ensembles. All experiments use a strict chemotype cluster split in which held-out test chemotypes are absent from the labelled and acquisition pools, providing a demanding evaluation of generalization to structurally distinct molecules.

Our results show that active-learning performance depends strongly on the endpoint, acquisition strategy, and uncertainty backend. Deep Ensembles frequently improve predictive performance relative to MC Dropout, but these gains do not consistently translate into better acquisition. Post-hoc analyses associate this disconnect with weak uncertainty-error alignment and reduced chemotype coverage in non-random batches. Diversity-aware acquisition produces favorable point estimates for several endpoints, motivating targeted replication, while random acquisition remains a competitive reference across the panel.

Together, these findings clarify when predictive uncertainty can support compound acquisition and highlight the importance of constructing informative, chemically diverse batches. With this work we provide a practical basis for selecting and validating acquisition strategies for individual ADMET endpoints and aim to motivate methods that better integrate uncertainty with chemotype coverage.

## Related Work

### Molecular Property Prediction

Classical ADMET models paired hand-crafted molecular descriptors (e.g., molecular weight, polar surface area, and count of hydrogen-bond donors) with random forests or support-vector machines ^9,10^. Graph neural networks (GNNs) moved beyond fixed descriptors by learning representations directly from the molecular graph, with PotentialNet ^4^ demonstrating large accuracy gains across several ADMET endpoints. Community benchmarks such as MoleculeNet ^31^ and the Therapeutics Data Commons (TDC) ^32^ subsequently enabled systematic comparisons across dozens of tasks and standardized evaluation protocols. Most of these models are trained on a fixed, randomly split dataset; how to select which molecules to label in order to build training sets efficiently has received comparatively little attention in the ADMET modeling literature, with a small number of exceptions discussed below ^33–35^.

### Uncertainty Quantification

Two practical families dominate current work, and we adopt both as the study’s uncertainty backends. **MC Dropout** ^36^ keeps dropout active at inference time and runs *T* stochastic forward passes; a heteroscedastic output head ^22^ supplies the epistemic–aleatoric decomposition at the cost of a single stored model. **Deep Ensembles** ^37^ train *M* independently initialized networks whose disagreement covers a broader region of parameter space ^38^, yielding better-calibrated intervals under distributional shift ^39^—a pattern confirmed on molecular property tasks ^40,41^—with *M* models to train and retrain per round. Wall-clock cost also depends on inference and acquisition. This cost asymmetry is why the present comparison is protocol-matched rather than compute-matched.

### Conformal Prediction

Mondrian conformal calibration ^42,43^ stratifies the calibration set on a grouping variable—here, binned predicted uncertainty. Under exchangeability, conformal methods can provide formal marginal coverage guarantees. In this benchmark, validation and test sets contain different held-out chemotype clusters, so that assumption need not apply. We therefore call the reported test intervals empirically calibrated intervals and evaluate their coverage empirically rather than claiming a formal guarantee under chemotype shift.

### Active Learning for ADMET

Pool-based uncertainty sampling ^16^ is the simplest AL strategy: each round, query the single unlabeled point about which the model is most uncertain. Chen *et al.* ^33^ applied this idea to site-of-metabolism prediction, identifying which atom in a drug molecule is modified by liver enzymes, and matched a fully supervised model while labeling only ~20% of atoms. Masood *et al.* ^34^ combined a pretrained molecular language model with a Bayesian AL loop for multi-task toxicity prediction, halving the labeling budget needed to reach a target accuracy. Bailey *et al.* ^35^ proposed COVDROP, which builds an epistemic covariance matrix from stochastic prediction samples and greedily maximizes joint batch variance via Cholesky decomposition. On permeability, solubility, and lipophilicity, COVDROP reached within 10% of the full-data RMSE with only a few hundred labeled molecules. Closest in intent to the present study, Yin *et al.* ^21^ benchmarked several uncertainty estimators for aqueous solubility and redox potential and then asked whether uncertainty-weighted selection accelerates generalization. They report improvement over a matched random batch for molecules outside the training distribution but none for in-distribution molecules, with the largest gains from an embedding-density criterion rather than a purely uncertainty-based one.

Each of these studies varies one axis while holding fixed something this benchmark varies. Yin *et al.* sweep the uncertainty estimator but perform a single acquisition step and score candidates individually rather than constructing the batch jointly; the remaining studies run iterative rounds but evaluate at most two acquisition strategies on one or two endpoints. No prior work compares batch-*construction* strategies—uncertainty-only, covariance-based, and diversity-aware—across a broad range of ADMET assay types over repeated rounds, nor examines how the choice of uncertainty backend (MC dropout vs. deep ensembles) interacts with acquisition strategy.

That gap matters more given how the method has fared under controlled scrutiny in other domains. Reevaluations in language ^19^ and vision ^20^ report that acquisition gains over random sampling are model- and task-dependent, sometimes vanish once stochasticity and regularization are held fixed, and are frequently not reproducible. Those studies motivate the two design commitments we adopt throughout: a random baseline matched on budget, split, and seed for every comparison, and uncertainty assessed across paired seeds alongside the learning curves.

### Batch Acquisition Functions

In practice, assays are run in plates of tens to hundreds of compounds, so selecting one molecule per AL round is unrealistic. Naive top-*k* uncertainty sampling simply picks the *k* most uncertain molecules, which often cluster in a narrow region of chemical space and provide redundant information.

Diversity-based methods spread the batch across the feature space. Core-set ^44^ selects points that minimize the maximum distance from any unlabeled molecule to its nearest labeled neighbor, ensuring broad coverage. BADGE ^45^ first computes “gradient embeddings”, vectors that capture both the model’s uncertainty and the direction of steepest loss, and then applies *k*-means++ to pick a diverse, high-uncertainty batch.

Covariance-based methods model the statistical dependencies within a candidate batch. Original BatchBALD ^46^ maximizes the mutual information between the joint labels of the batch and the model parameters, preferring batches whose members are individually uncertain yet complementary. COVDROP ^35^ constructs an epistemic covariance matrix from a set of stochastic predictions: each entry encodes how correlated two molecules’ prediction uncertainties are. A greedy Cholesky decomposition then selects molecules one by one, at each step choosing the candidate that adds the most independent variance to the batch, effectively avoiding redundant selections without an explicit distance computation. We call our noise-normalized covariance variant **Noise-Normalized Covariance Acquisition (NN-Cov)**. NN-Cov is inspired by BatchBALD’s information-gain motivation but is not the original joint-mutual-information algorithm. For reproducibility, released directories and machine-readable artifacts retain the legacy identifier batchbald; no artifact key has been renamed.

## Materials and Methods

We now describe the full experimental pipeline used to generate the results in this study—from dataset curation and chemotype-aware splitting, through the Bayesian GNN and its two uncertainty backends, to the seven acquisition strategies and the paired statistical evaluation that determines whether non-random selection actually outperforms random sampling. Figure 1 gives a schematic overview of how these components connect end to end, before each is described in full detail in the subsections that follow.

**Figure 1:**
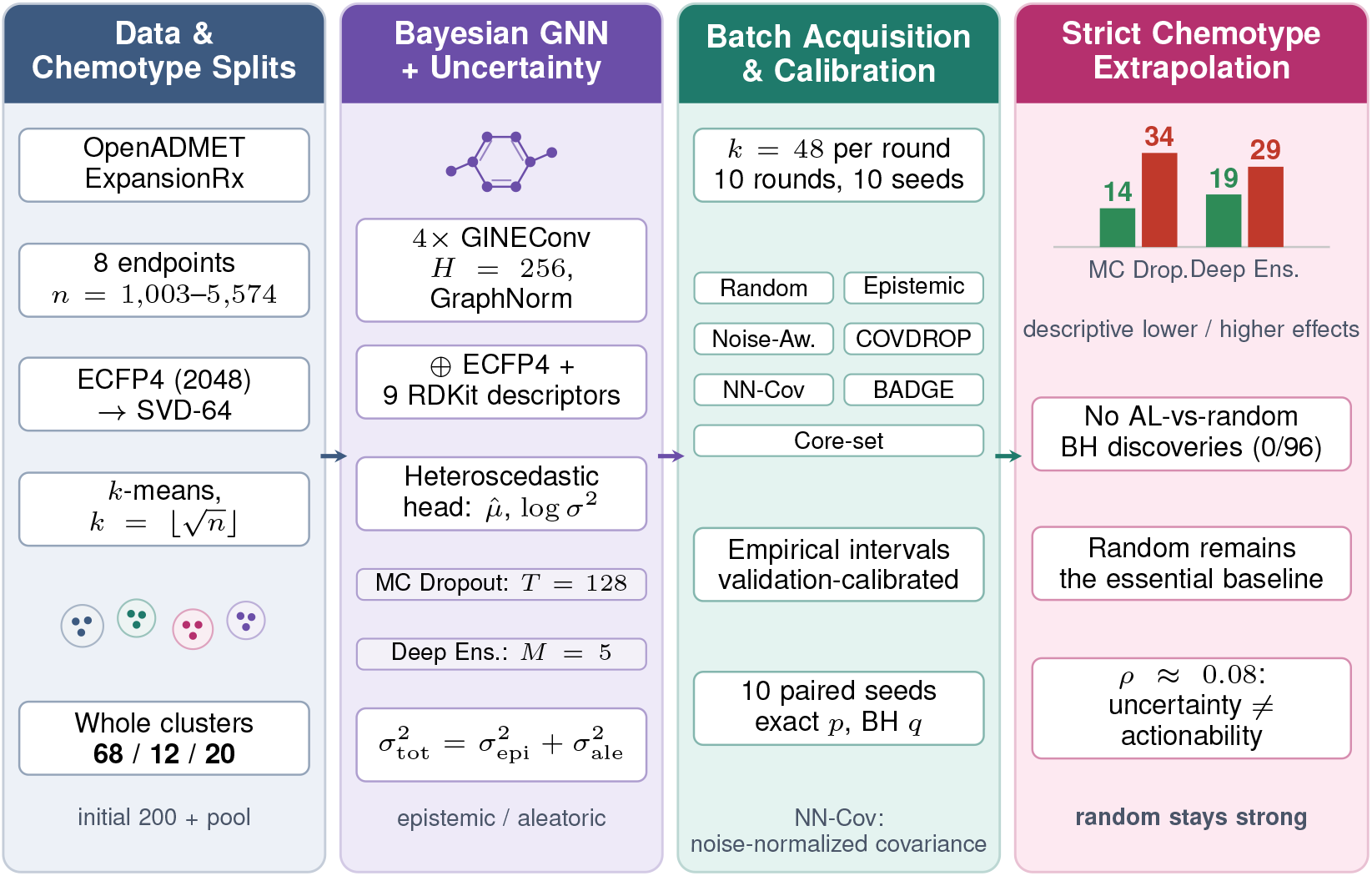
End-to-end overview of the benchmark. Molecules from the OpenADMET ExpansionRx dataset are clustered by ECFP4 similarity and whole chemotype clusters are assigned to a fixed 68/12/20 split (left). A Bayesian GNN fusing graph, fingerprint, and descriptor features predicts each endpoint through a heteroscedastic head, with predictive variance decomposed into epistemic and aleatoric components under two backends (center left). Seven acquisition strategies each select *k* = 48 molecules per round over ten rounds and ten seeds, with Mondrian conformal calibration and paired seed-level inference against a matched random baseline (center right). The resulting comparison shows endpoint-dependent effects and an uncertainty–actionability gap (right).

### Exploratory Data Analysis

We use the OpenADMET ExpansionRx dataset from Hugging Face ^30^, a curated collection of ADMET measurements for drug-like small molecules. Each endpoint is an independent regression task; molecules with missing labels for a given endpoint are excluded, yielding between 1,003 (MBPB) and 5,574 (KSOL) examples per task. The eight endpoints span four pharmacological property classes ^1^:

- **Physicochemical:** LogD (lipophilicity) and KSOL (kinetic aqueous solubility).
- **Metabolic stability:** HLM CLint and MLM CLint (intrinsic clearance in human and mouse liver microsomes, respectively).
- **Intestinal transport:** Caco-2 Papp A→B (apparent permeability through a model intestinal cell layer) and Caco-2 Efflux Ratio (efflux transporter activity).
- **Murine protein binding:** MPPB and MBPB (fraction of compound unbound in mouse plasma or brain; only the unbound fraction is pharmacologically active).

**Summary statistics** and target-transform choices are shown in Table 1. LogD is broadly symmetric (skew = −0.30) and uses the identity target transform. Metabolic clearance spans several orders of magnitude and includes zero-valued measurements. The Caco-2 efflux ratio has a median near unity (1.38) but reaches 105.64, indicating that a subset of compounds is actively pumped out by intestinal transporters. Protein binding fractions are concentrated at low unbound values, meaning most compounds are heavily bound to plasma or tissue proteins.

**Table 1:**
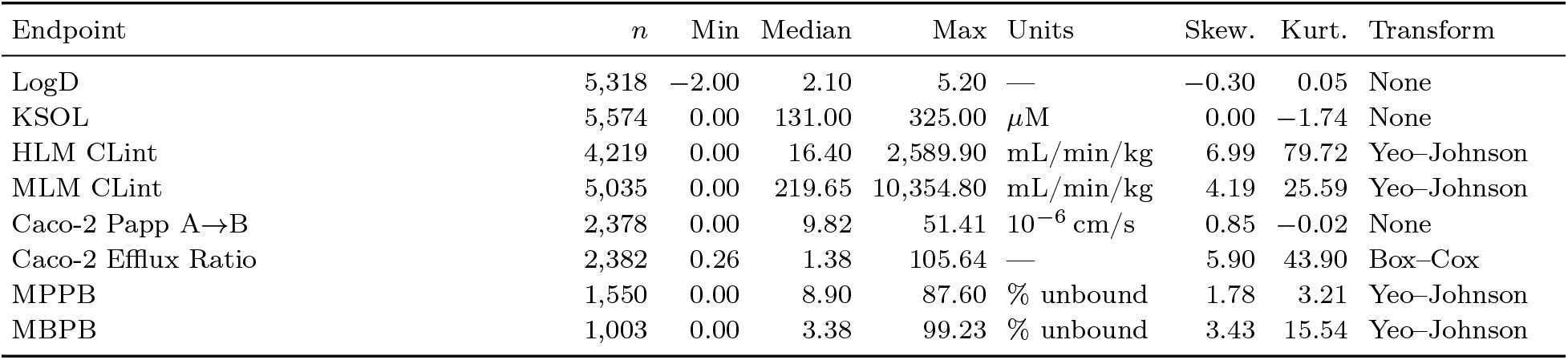
Endpoint summary statistics and target transformations.

**Target transformations.** Strongly skewed targets, as in clearance and protein-binding endpoints, can complicate regression and increase the influence of extreme values during training. We therefore select a variance-stabilizing transform per endpoint based on skewness and kurtosis (Table 1). LogD, KSOL, and Caco-2 Papp are near-symmetric and require no transformation. The two clearance endpoints and two protein binding endpoints are right-skewed; we apply the Yeo–Johnson transform ^47^, a flexible power transform that, unlike Box-Cox, handles zero and negative values. Caco-2 Efflux Ratio is strictly positive and similarly skewed; Box–Cox is used, with an additive shift if needed to ensure positivity. All models train on the endpoint-specific target scale. RMSE, MAE, and interval widths are reported in original target units, and coverage is assessed against original-unit labels. Gaussian NLL is evaluated on the transformed scale using the predictive mean and total variance; it is compared within matched endpoints.

**Cluster-based splitting.** A random train/validation/test split is problematic for molecular data: structurally similar molecules (within the same chemotype or scaffold family) may land on both sides of the split, allowing the model to generalize superficially by memorizing local structure-activity relationships rather than learning transferable rules.

To evaluate transfer between chemical groups, we use a strict chemotype cluster split: molecules are clustered by structural similarity, and entire clusters are assigned to a single partition. This design measures generalization to held-out clusters; its relevance to a prospective campaign depends on the overlap between available and deployment chemistry. Because each endpoint is an independent regression task with its own set of labeled molecules (Table 1), clustering and splitting are performed separately per endpoint. For a given endpoint with *n* unique featurizable molecules, each molecule is encoded as an ECFP4 circular fingerprint (2,048 bits, radius 2), dimensionality is reduced to 64 with truncated SVD, and clustering is performed with MiniBatchKMeans using 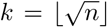 clusters, clipped to [10, 200]. Here *n* is the deduplicated, featurizable population for that endpoint rather than the raw labeled-row count of Table 1; both values, and the resulting *k*, are tabulated per endpoint in Supplementary Table S3 so the rule is directly checkable. The resulting cluster-size distribution has a long tail: a few dominant chemotypes account for most molecules, while many clusters are sparsely populated (Fig. 2b). A UMAP projection ^48^ confirms this heterogeneity (Fig. 2a).

**Figure 2:**
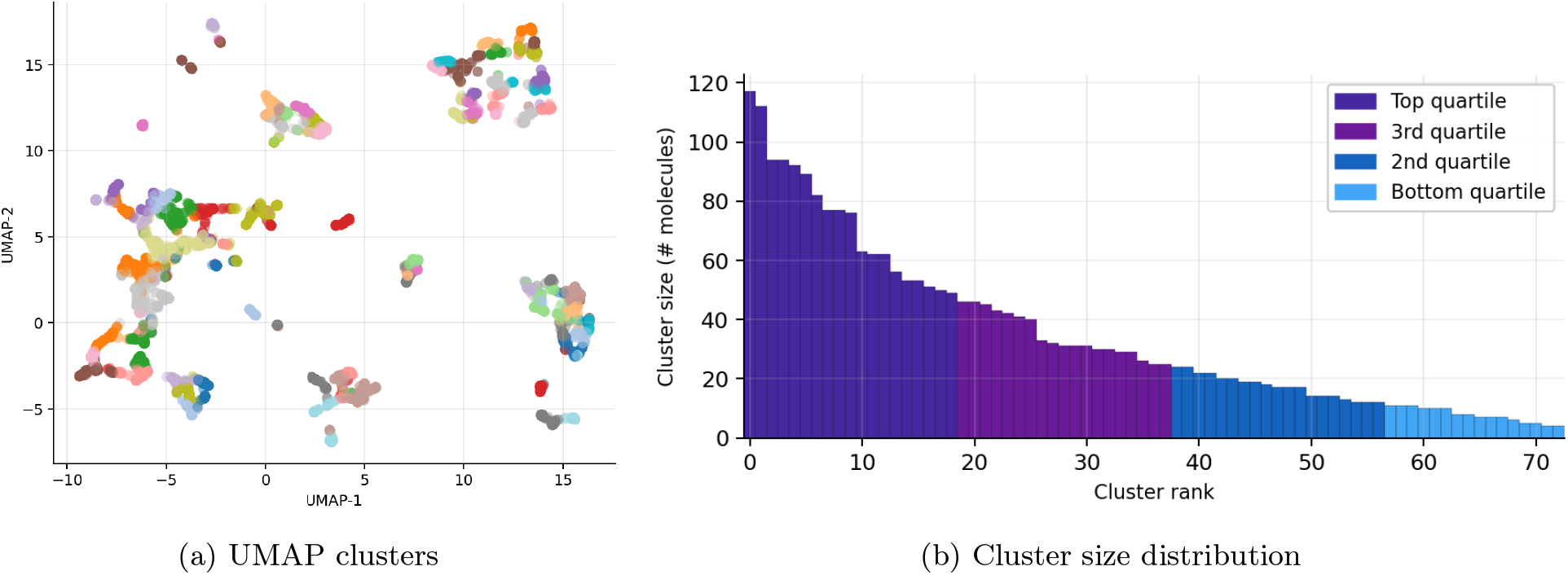
UMAP visualization and cluster size distribution.

We assign whole clusters to train/validation/test in a 68/12/20 ratio. The 20% test set provides stable metric estimation; the 12% validation set supports early stopping and Mondrian conformal calibration; and the remaining 68% forms the training pool. Within that train pool, the initial 200 molecules are selected by a seeded stratified shuffle over 12 quantile-style bins of the *train-pool target labels*; the remainder forms the unlabeled acquisition pool. Thus initialization is target-informed. No validation or test labels enter this construction. Table 1 reports raw labeled-row counts, whereas the split audit reports the endpoint-specific featurizable population and counts clusters using unique SMILES; SMILES-level de-duplication and dropped unfeaturizable molecules therefore make the raw and split counts differ, by between 3% (MBPB) and 16% (MPPB). Supplementary Table S3 gives both counts side by side. Because *n* varies across endpoints, the number of clusters and the absolute partition sizes differ per endpoint, while the 68/12/20 proportion is held fixed throughout. This chemotype-level separation is maintained throughout all experiments. Consequently, held-out test clusters never appear in the acquisition pool: no strategy can query molecules from those test chemotypes, and reported metrics measure generalization to held-out chemical groups.

### Model Architecture

Given the heterogeneity of ADMET endpoints described above, we design a model that fuses graph, fingerprint, and descriptor features to capture information at different scales.

Each molecule is first converted into a graph where atoms are nodes (featurized by element type, formal charge, hybridization, etc.) and bonds are edges (featurized by bond type, stereochemistry, and ring membership). Four GINEConv message-passing layers (*H* = 256 hidden channels) propagate information across the graph. GINEConv ^49^ extends the Graph Isomorphism Network ^50^, which is maximally expressive among message-passing architectures under the Weisfeiler–Lehman test, so that edge features participate in aggregation; this matters here because bond order and ring membership carry much of the signal for permeability and metabolic stability. Each layer is followed by GraphNorm ^51^, chosen over batch normalization because graph-level batch statistics are noisy at the small batch sizes an active-learning loop operates under, together with virtual-node augmentation, residual connections, and dropout (*p* = 0.20). A global attention pooling layer aggregates the result into a graph-level vector **g** ∈ ℝ^256^.

To complement these learned features, we concatenate ECFP4 fingerprints (2,048 bits) with nine physicochemical descriptors computed with RDKit ^52^ (e.g., molecular weight, topological polar surface area, number of rotatable bonds), yielding an auxiliary vector **a** ∈ ℝ^2057^. Retaining fixed descriptors alongside a learned representation is a deliberate hedge, since the two views degrade differently on unseen chemotypes ^7^.

The graph and auxiliary vectors are concatenated into **h** = [**g** || **a**] ∈ ℝ^2313^ and fed into two parallel prediction heads. A mean head (Linear→ReLU→Dropout→Linear(1)) produces the point estimate *µ̂*, and a log-variance head of identical architecture (output clamped to [−10, 5]) predicts the per-molecule aleatoric noise 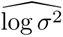. The model is trained by minimizing the Gaussian negative log-likelihood (NLL). Figure 3 shows how this shared architecture is instantiated under each uncertainty backend. Both panels use the identical GNN trunk and the two parallel prediction heads described above; they differ only in how the *S* stochastic samples are generated, by *M* independently trained ensemble members in panel (a) and by *T* dropout-active forward passes through a single model in panel (b). The variance decomposition of Eq. (1) is applied identically in both cases.

**Figure 3:**
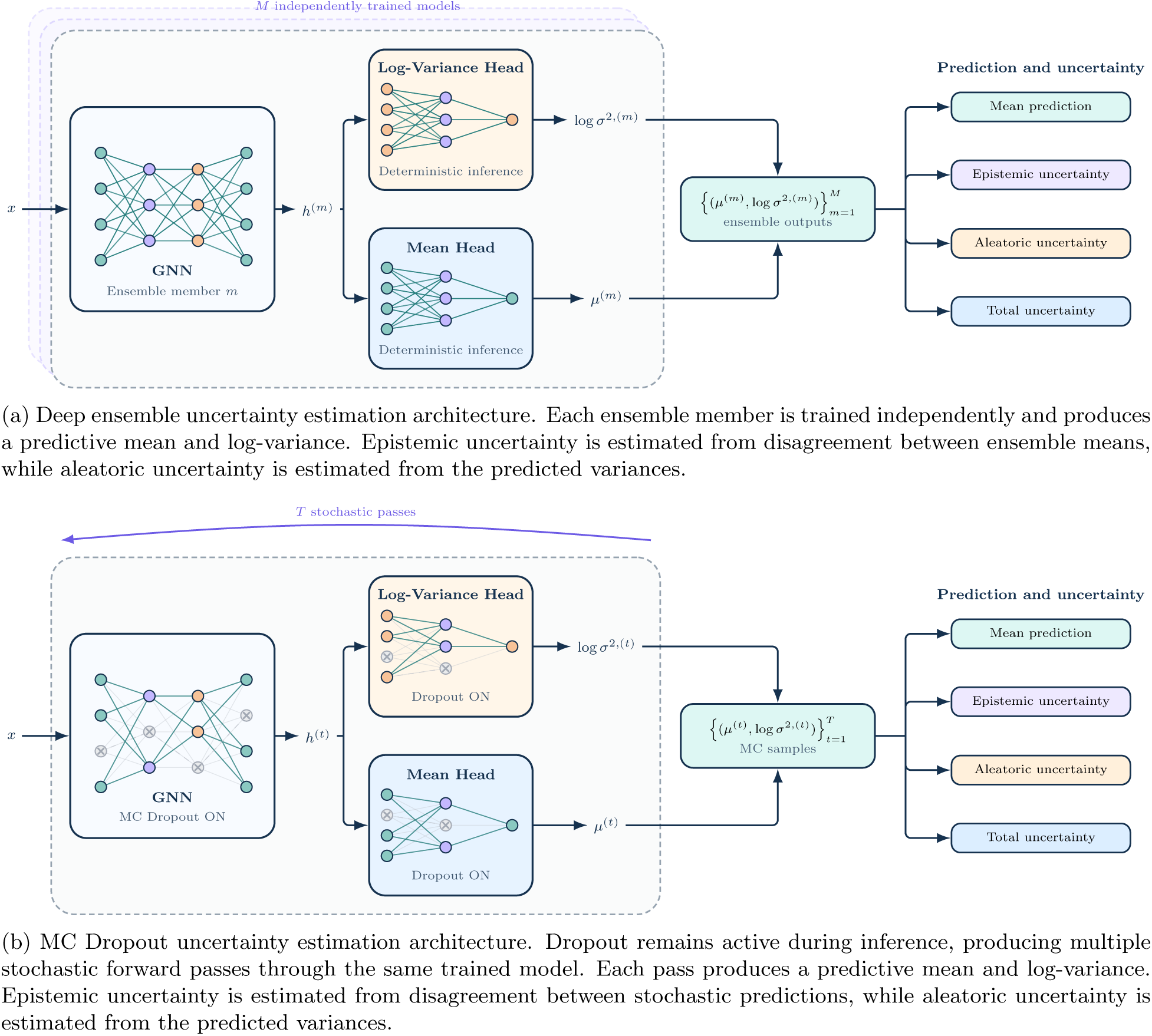
Matched uncertainty-estimation architectures used in the benchmark. Panel (a) shows the deterministic deep-ensemble backend; panel (b) shows stochastic MC-Dropout inference. Both use the same heteroscedastic prediction heads and variance decomposition.

**Figure 4:**
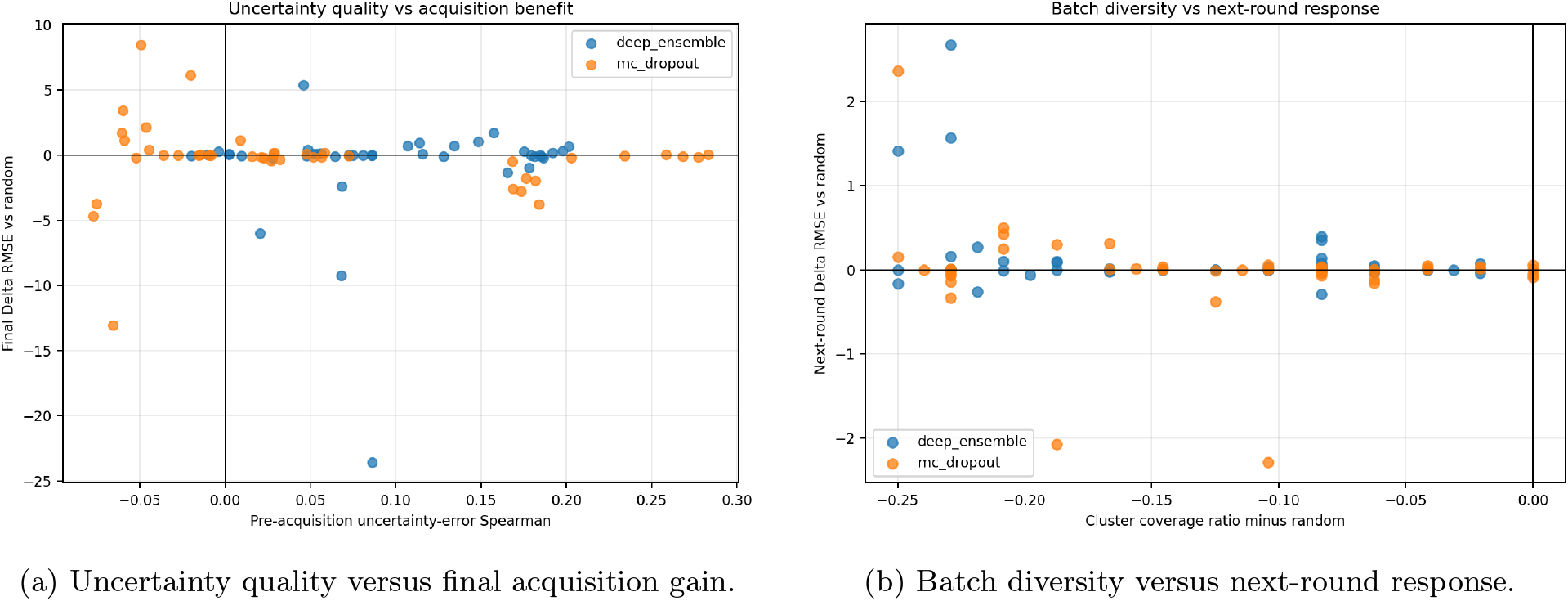
Post-hoc actionability diagnostics. Panel (a) compares pre-acquisition uncertainty–error ranking with final ΔRMSE versus random. Panel (b) compares selected-batch cluster coverage against next-round ΔRMSE versus random. Negative values indicate lower final RMSE in panel (a) or a more favorable round-to-round RMSE change in panel (b), relative to random.

### Fixed-Data Classical Baseline

As a predictor-competitiveness check, we fit a Random Forest to the exact final labeled set produced by each seed’s random-acquisition trajectory. This is a fixed-data comparison, not a third uncertainty backend and not a Random-Forest active-learning experiment. The representation comprises a chiral 2,048-bit ECFP4 fingerprint (radius 2) and the same nine RDKit descriptors used by the GNN auxiliary block. Median imputation is fit only on the labeled training set. The prespecified Random Forest uses 500 trees, bootstrap sampling, `max_features=sqrt`, and `min_samples_leaf=1`; validation metrics are diagnostic only, with no hyperparameter or model selection, and test data are used only for final evaluation. Each fit is paired to both GNN random baselines on identical test molecules and labels at the same final budget. Round 10 is used except for the documented MBPB seeds 0, 5, and 6, which use their final common round 9.

### Uncertainty Estimation

Regardless of backend, we decompose total predictive variance into two components. Let *µ̂*, denote the mean prediction from the *s*-th stochastic sample (a dropout pass or an ensemble member). Given *S* such samples,

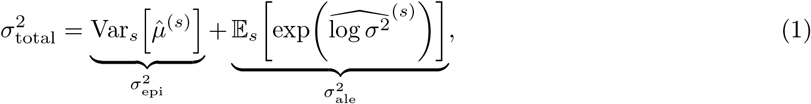

where 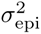 captures variance arising from limited training data and 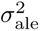 reflects irreducible measurement noise. The predictive mean is 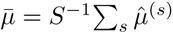. We also track the mutual-information proxy 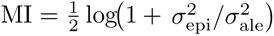 : a high MI indicates greater epistemic uncertainty relative to predicted aleatoric noise; whether querying these molecules reduces error is evaluated empirically.

The two backends differ in how *S* is generated and in their cost–quality trade-off.

#### MC Dropout

MC dropout ^36^ keeps all dropout layers ^53^ active at inference time and performs *T* stochastic forward passes (*S* = *T*).

Running *T* passes over a large unlabeled pool at every AL round is expensive, so we adopt a two-stage strategy: a rapid screen with *T*_screen_ = 8 identifies a shortlist (top 10 × *k* candidates by epistemic uncertainty), and a refined evaluation with *T*_acq_ = 96 scores only the shortlist. For calibration and final evaluation we use *T*_val_ = *T*_test_ = 128.

#### Deep Ensembles

We train *M* = 5 independently initialized members ^37^, following the convention in molecular property prediction ^41^. At inference every member runs with dropout disabled, yielding *S* = *M* samples. No two-stage screening is needed since all acquisition functions operate on full-pool ensemble predictions directly; for covariance-based methods (COVDROP, NN-Cov) we still shortlist by epistemic uncertainty to keep the *N × N* matrix tractable.

Both COVDROP and NN-Cov construct their covariance matrices from a generic set of stochastic predictions—whether *T* dropout passes or *M* ensemble members—so the Cholesky batch-selection logic is identical in both cases. NN-Cov uses the tractable normalization described below because evaluating full joint label–parameter mutual information over the molecular pool is not computationally practical at benchmark scale.

### Training and Fine-Tuning

All models are optimized with AdamW ^54^ (*η* = 3×10^−4^, *λ* = 2×10^−4^) under a schedule of 12% linear warmup followed by cosine decay, with gradient norms clipped at 1.0. Early stopping monitors validation NLL rather than validation RMSE, so that the log-variance head must converge alongside the mean head before training halts. Patience is 18 for the initial fit and is reduced to 9 for the fine-tuning rounds, which begin from an already-trained checkpoint.

The two backends differ only in how this recipe is instantiated. Under the **MC dropout backend**, a single model is trained on the endpoint’s protocol-defined initial labeled set of 200 molecules, then *warm-started* from the previous round’s checkpoint at each AL round with a fresh optimizer. Under the **ensemble backend**, each of the *M* = 5 members is independently initialized with a distinct random seed and trained from scratch on the same initial set; during AL rounds every member is warm-started from its own checkpoint and fine-tuned, again with a fresh optimizer. Hyperparameters are identical across members and across backends, which is what makes the backend comparison protocol-matched rather than separately tuned.

### Mondrian Conformal Calibration

For every trained model, Mondrian conformal calibration constructs empirically calibrated prediction intervals ^42,43,55^.

Calibration uses held-out validation labels and predictions only. For the seven endpoints other than Caco-2 Efflux Ratio, validation molecules are grouped by quantiles of predicted total standard deviation on the transformed scale, using up to four bins with at least 12 molecules per bin. Within each bin, a scale *c_b_* is fitted by bisection over [10^−3^, 32] to target 0.95 empirical coverage after inverse transformation. The interval for a prediction *µ̂_t_* with standard deviation *σ̂_t_* in bin *b* is 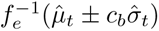. If the requested bins are too small, MC Dropout falls back to a global scale; Deep Ensemble first tries fewer bins before using global calibration.

Caco-2 Efflux Ratio uses the locked *c*–*τ* calibration variant. In each valid uncertainty bin, finite-sample-adjusted quantiles of transformed-scale absolute residuals divided by predicted standard deviation determine *c_b_*; a corresponding residual quantile determines the additional scale *τ_b_*. Both scales receive the prespecified safety multiplier of 1.10. The transformed interval half-width is 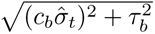, and the bounds are then inverse-transformed. If any bin has fewer than 12 molecules, a global *c*–*τ* calibration is used. This variant is shared by both backends and does not use the bisection bound of 32.

Calibration is recomputed after every AL round. Validation and test occupy disjoint held-out chemotype clusters, so standard exchangeability need not hold; the resulting test intervals are reported as empirically calibrated intervals with observed coverage, not as intervals carrying an unqualified formal conformal guarantee.

### Batch Acquisition Strategies

Each AL round selects a batch of *k* = 48 molecules from the unlabeled pool for simulated measurement. For the eight endpoints in the final paired analysis, we run up to 10 rounds from each endpoint’s protocol-defined initial labeled set. This retrospective initial set is selected using train-pool target labels through 12-bin regression stratification, reducing target-range imbalance before acquisition; no validation or test labels are used. This construction is not directly realizable in a prospective campaign whose pool labels are genuinely unknown. We repeat every experiment over 10 random seeds. The primary estimand is the mean of the ten paired seed-level metric differences. We use *B* = 10,000 paired-seed bootstrap resamples for a 95% confidence interval, an exact two-sided paired sign-flip test for the raw p-value, and Benjamini–Hochberg correction within prespecified families at *α* = 0.05. Bootstrap resampling estimates uncertainty over ten paired seed-level differences; *B* = 10,000 controls Monte Carlo precision and does not increase the effective number of experimental replicates. The acquisition functions below use the same definitions under both uncertainty backends. Non-random methods use uncertainty estimates, prediction covariances, or embeddings as specified below; random acquisition is independent of the uncertainty backend.

1. **Random.** Uniform sampling from the unlabeled pool; the matched reference for evaluating acquisition gains.
2. **Epistemic Top-***k*. Select the *k* molecules with the highest 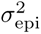. Under MC dropout, the two-stage screen described under MC Dropout narrows the pool first; under ensembles, the full pool is scored directly.
3. **Noise-Aware.** Rank by 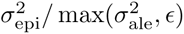 with *ɛ* = 10^−8^, down-weighting molecules whose uncertainty is dominated by irreducible assay noise while preventing numerical instability when the aleatoric estimate is very small.
4. **COVDROP** ^35^. Builds an epistemic covariance matrix from the *S* stochastic predictions and applies greedy Cholesky decomposition to select the batch that maximizes joint independent variance.
5. **Noise-Normalized Covariance Acquisition (NN-Cov)** ^46^. Same greedy Cholesky procedure as COVDROP, but on a noise-normalized covariance (epistemic divided by aleatoric) to prevent high-noise molecules from dominating selection. This is the tractable covariance-based variant described above, not the original joint-mutual-information algorithm.
6. **BADGE** ^45^. Applies *k*-means++ to last-layer gradient embeddings scaled by *σ*_epi_.
7. **Core-set** *k***-center**^44^. Greedy *k*-center in GNN embedding space **g**, weighted by epistemic uncertainty.

## Results and Discussion

The benchmark examines three complementary aspects of ADMET active learning: endpoint-specific acquisition effects, the choice of predictive backend, and the relationship between uncertainty and batch composition. The final paired seed-level analysis uses eight validated endpoints (Supplementary Table S2): LogD, KSOL, HLM CLint, MLM CLint, Caco-2 Papp A→B, Caco-2 Efflux Ratio, MPPB, and MBPB. All results use the strict chemotype cluster split. For MBPB, seeds 0, 5, and 6 are missing matched round-10 outputs across all acquisition methods and both backends; paired analyses therefore use the final common available round for those matched comparisons.

We report final-round active-learning performance relative to the random acquisition baseline. Negative ΔRMSE means lower mean RMSE than random under the same backend, endpoint, seed, and labeling budget; positive values mean worse RMSE. The table reports effect sizes descriptively; full seed-bootstrap intervals, exact raw p-values, BH q-values, and paired seed differences are provided in the Supplementary Materials and machine-readable artifacts.

### Active Learning Is Endpoint-Dependent

Table 2 shows the endpoint dependence of acquisition effects. Mean final RMSE is lower than matched random in 14/48 MC-Dropout and 19/48 Deep-Ensemble method–endpoint cells, and higher in 34/48 and 29/48, respectively. KSOL has the broadest favorable pattern, while other endpoints show smaller or method-specific differences. These descriptive effects help identify candidates for replication; they do not establish a uniformly beneficial acquisition strategy.

**Table 2:**
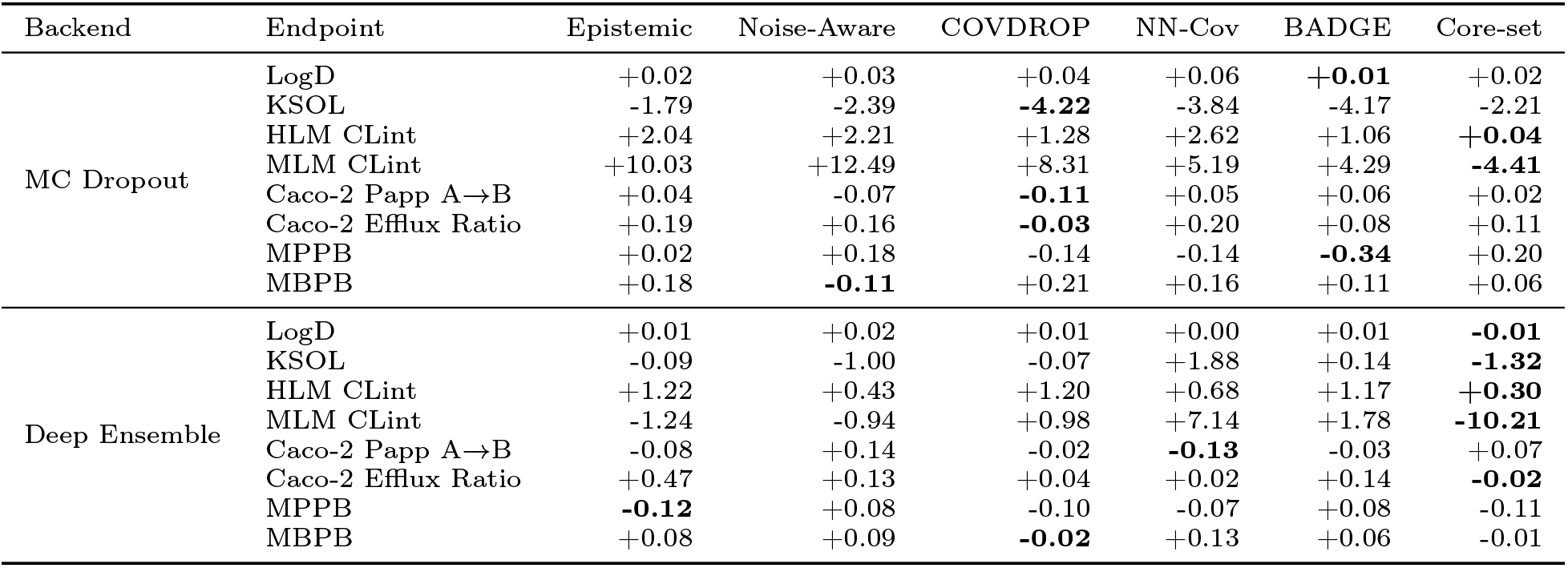
Mean final-round paired-seed ΔRMSE versus random. Negative values indicate descriptively lower RMSE than random; bold marks the best non-random point estimate per endpoint and backend. Seed-bootstrap intervals and exact raw p-values and BH q-values are reported in the Supplementary Materials.

**Table 3:** Backend-level summary. Negative/positive effects are descriptive mean paired-seed ΔRMSE counts over 48 acquisition-versus-random cells. Raw rejections use exact two-sided sign-flip *p <* 0.05; adjusted discoveries use BH *q <* 0.05 within each backend’s 48-test RMSE family.

| Backend | Endpoints | Negative/positive effects | Raw rejects | BH discoveries |
| --- | --- | --- | --- | --- |
| MC Dropout | 8 | 14/34 | 5/48 | 0/48 |
| Deep Ensemble | 8 | 19/29 | 2/48 | 0/48 |

### Acquisition Opportunities Across Endpoints

KSOL has the broadest descriptive acquisition gains: all six MC-Dropout point estimates are negative, as are four of six Deep-Ensemble estimates. The largest are MC-Dropout COVDROP (ΔRMSE = −4.22) and BADGE (−4.17), and Deep-Ensemble Core-set (−1.32). Caco-2 Papp A→B and MPPB also contain selected negative point estimates, while Core-set has favorable estimates for MLM CLint under both backends. These patterns nominate endpoint–method combinations for replication, with paired uncertainty estimates available in the Supplementary Materials.

### Endpoints Where Random Acquisition Remains Competitive

Random acquisition remains competitive on several endpoints. All six MC-Dropout LogD effects are positive, although small; five of six Deep-Ensemble effects are also nonnegative. HLM CLint has positive effects across all methods and both backends, with Core-set closest to random. MLM CLint, Efflux, MPPB, and MBPB show mixtures of positive and negative effects. Together with the favorable KSOL pattern, these results indicate that acquisition choices should reflect endpoint-specific behavior and the magnitude and variability of paired effects.

Under the strict chemotype cluster split, acquisition can only improve representations using train-pool chemotypes; it cannot label the held-out test clusters themselves. The descriptive pattern suggests that acquisition is more promising when uncertainty aligns with reducible model error and the selected batch adds chemical diversity, and less promising when random acquisition already covers the available pool well, assay noise dominates, or high-uncertainty molecules form redundant pockets. These post-hoc hypotheses and conclusions are confined to the evaluated extrapolation regime.

### Diversity-Aware Acquisition Patterns

Acquisition-family rankings remain descriptive. Core-set has several favorable point estimates, including Deep-Ensemble KSOL and MLM CLint, while COVDROP and BADGE produce large negative point estimates in selected cells. Pure Epistemic Top-*k* and Noise-Aware sampling have more positive than negative RMSE effects under MC Dropout. The variation within and between families motivates evaluating chemical-space coverage alongside uncertainty ranking, without identifying any family as uniformly superior.

This descriptive pattern is consistent with chemical-space coverage mattering alongside uncertainty ranking, but does not establish their relative causal effects. Diversity-aware acquisition can prevent the model from spending an entire batch on near-duplicate high-uncertainty compounds, a failure mode that is especially plausible in clustered molecular libraries. The same ordering appears in the external evaluation closest to ours, where an embedding-density criterion outperformed purely uncertainty-based selection on molecular property prediction ^21^; this provides a complementary motivation for testing diversity-aware criteria across endpoints and models.

### MC Dropout Versus Deep Ensembles

The matched backend comparison helps distinguish predictor quality from acquisition effects. Deep Ensembles have lower mean RMSE in most endpoint–method cells, while the value of non-random selection remains endpoint-dependent.

Across 56 matched endpoint–method cells, Deep Ensemble has lower mean RMSE in 45 and higher mean RMSE in 11. The clearest supported differences favor Deep Ensemble on six KSOL cells (all methods except NN-Cov) and NN-Cov on Caco-2 Papp A→B. These effects range from 6.08 to 10.40 RMSE units for KSOL (95% seed CIs all below zero) and 0.447 for Papp NN-Cov (95% CI [ 0.770, 0.208]). The paired analysis also supports Deep Ensemble in a subset of MAE and NLL comparisons, while ACE95 does not establish a consistent calibration advantage. Full raw and adjusted inference is reported in the Supplementary Materials.

This separation matters for interpreting the study. Deep Ensembles train exactly 5× as many models as MC Dropout, which follows by construction from *M* = 5 rather than from measurement. Measured wall-clock cost does not follow that ratio, because MC Dropout incurs substantially more stochastic inference per trained model. In Supplementary Table S10 the MC Dropout range (10.10–14.71 h across acquisition methods) straddles the much narrower Deep Ensemble range (11.19–12.22 h): the cheapest MC Dropout methods are faster than any Deep Ensemble configuration, while the most expensive run about 20% longer than Deep Ensemble’s slowest (14.71 versus 12.22 h). Wall-clock cost is therefore driven more by the choice of acquisition function under MC Dropout than by the choice of backend, and a single “comparable” or “cheaper” verdict would misrepresent the table. The acquisition problem nevertheless remains endpoint-dependent. Predictive accuracy and acquisition benefit should therefore be assessed separately for each ADMET endpoint.

### The Fixed-Data Random Forest Is Endpoint- and Metric-Dependent

The classical baseline provides a deliberately narrower test of predictor competitiveness at the same labeled budget. Across the 16 endpoint–GNN-backend comparisons per metric, BH-adjusted RMSE results favor the Random Forest in 5 cells, a GNN in 6, and neither in 5; adjusted MAE favors the Random Forest in 1 cell, a GNN in 13, and neither in 2 (Table 4). The RMSE results favor the Random Forest against both GNN backends on Caco-2 Papp A→B and HLM CLint, and against MC Dropout on KSOL. They favor both GNNs on LogD, MBPB, and MPPB. For MAE, only Caco-2 Papp A→B against MC Dropout favors the Random Forest after correction, while the GNNs are favored broadly. Full endpoint effects, seed-bootstrap intervals, exact paired sign-flip p-values, and BH q-values are reported in the Supplementary Materials.

**Table 4:** Adjusted fixed-data Random Forest versus GNN results. Each metric has 16 paired endpoint–backend comparisons. “RF” and “GNN” count the favored predictor among BH discoveries; “Neither” denotes no adjusted difference.

| Metric | RF favored | GNN favored | Neither |
| --- | --- | --- | --- |
| RMSE | 5 | 6 | 5 |
| MAE | 1 | 13 | 2 |

These results do not support a universal architecture ranking. RMSE and MAE emphasize different parts of the error distribution, and the direction changes across endpoints. The comparison also does not test whether Random-Forest uncertainty would improve acquisition: it holds the acquired data fixed and evaluates only the predictor fitted afterward.

### Uncertainty Quality Does Not Guarantee Acquisition Usefulness

To understand the variation in acquisition outcomes, we ran a post-hoc “uncertainty actionability” analysis over the existing final artifacts. This analysis uses test-set prediction errors only after the experiments are complete; test data was never used for acquisition, calibration, tuning, model selection, or run filtering. The goal is to separate two questions that are often conflated: whether uncertainty ranks molecules with high prediction error, and whether labeling the molecules selected by that uncertainty improves the next model. The post-hoc diagnostics are consistent with an uncertainty–actionability gap. Across non-random acquisition batches, the median pre-acquisition Spearman correlation between epistemic uncertainty and absolute test error is only 0.078. The top 10% most epistemically uncertain test molecules have only 1.14× the mean absolute error of the remaining molecules, and the overlap between the top uncertainty decile and top error decile is 0.111, only slightly above the 0.10 expected by chance. Thus, the observed uncertainty–error association is positive but weak. More importantly, even this signal does not automatically translate into useful labels: the median next-round ΔRMSE versus random is +0.0033, and the median final-round ΔRMSE versus random is +0.018 (both pooled across all matched non-random cells, in contrast to the per-method medians in Table 5). Positive values indicate a less favorable round-to-round change or higher final RMSE than random, respectively.

**Table 5:**
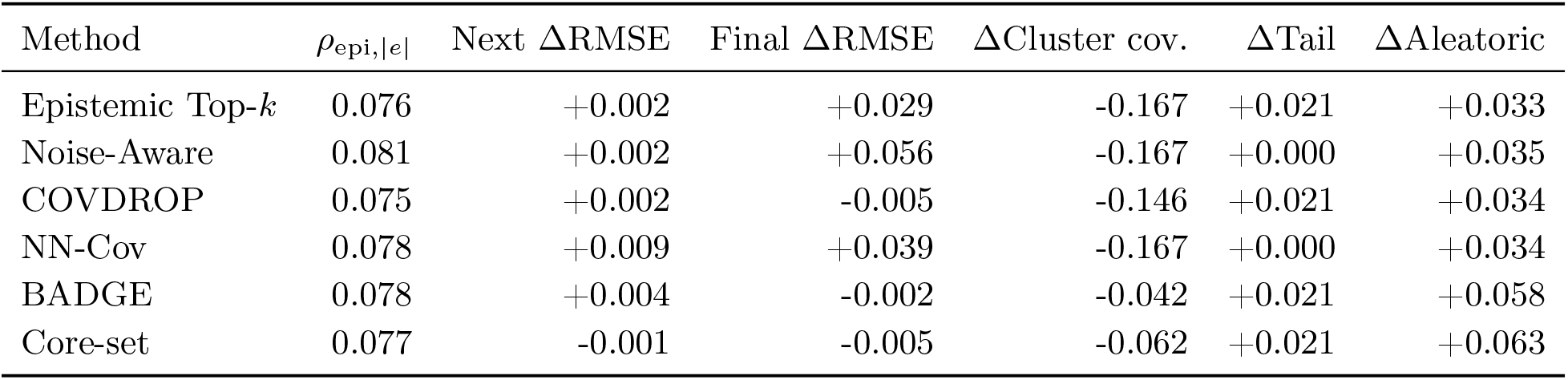
Post-hoc method-level actionability diagnostics. Values are medians over matched backend, end-point, seed, and round cells. Deltas are differences from the matched random trajectory or batch, not normalized ratios. RMSE differences retain each endpoint’s original units; pooled medians are descriptive summaries across heterogeneous scales. Column definitions: *ρ*_epi_,_|_*_e_*_|_ is the Spearman correlation between epistemic uncertainty and absolute test error; Next ΔRMSE is the method’s round-to-round RMSE change minus the matched random change; Final ΔRMSE is method minus random RMSE at the final common round; negative values favor the method; ΔCluster cov. is the cluster-coverage ratio minus that of the matched random batch; ΔTail is tail-label enrichment relative to random; and ΔAleatoric is aleatoric-fraction enrichment relative to random.

The backend comparison illustrates this distinction. Deep Ensembles rank high-error molecules better than MC Dropout by epistemic uncertainty (median pre-acquisition Spearman 0.092 versus 0.059), consistent with their stronger predictive performance, and their top-decile error enrichment is also slightly higher (1.15 versus 1.12). Yet Deep Ensemble acquisitions are still not uniformly useful: their median next-round and final-round ΔRMSE values versus random remain slightly positive (+0.0033 and +0.017). The two backends also give up exactly the same median cluster coverage relative to random (−0.125), indicating similar batch coverage despite the difference in uncertainty ranking. Supplementary Table S9 reports these diagnostics split by backend and by method, using the same cells and column definitions as Table 5. Stronger predictive performance therefore co-occurs with better uncertainty ranking here, but not with consistently better acquisition outcomes; this association does not establish a causal pathway.

Several factors could be associated with this uncertainty–actionability pattern. First, the heteroscedastic likelihood may not fully separate assay noise from representation or model misspecification, so a large predicted epistemic component need not identify reducible error. Second, warm-starting across rounds may preserve a narrow basin of solutions and make posterior disagreement overconfident or insufficiently diverse. Third, the cluster split evaluates transfer between held-out chemical groups: errors may reflect irreducible gaps between chemotypes rather than interpolation errors that additional labels from the selected region can repair. These are hypotheses rather than causal conclusions; the present post-hoc analysis cannot determine whether any of them produces the observed acquisition pattern.

The third hypothesis also offers a possible reconciliation with the reported gains of uncertainty-guided selection on molecules outside the training distribution^21^. The relationship between pool and evaluation chemistry may influence the value of acquisition. Here whole chemotype clusters are assigned to the test partition and are therefore absent from the pool by construction, so no acquisition rule—however well calibrated—can sample the chemistry on which it is ultimately scored. This difference in pool composition motivates comparing split regimes directly to determine when uncertainty-guided selection is most useful.

Our post-hoc analyses associate random acquisition’s competitiveness with batch composition. Compared with the random batch selected under the same backend, endpoint, seed, and round, non-random batches cover fewer train-pool clusters in median (Δ cluster coverage ratio = −0.125), show slightly higher tail-label enrichment (+0.0208), and have higher aleatoric-fraction enrichment (+0.0346). A coverage loss of 0.125 corresponds to roughly six fewer unique clusters in a 48-molecule batch. The common 200-molecule initial set may already cover much of the learnable space for some endpoints. Moreover, because it is selected using train-pool target labels through 12-bin regression stratification, it may be unusually informative and reduce the room for subsequent acquisition to improve; a 48-molecule random batch then provides substantial stochastic diversity. For comparatively easy endpoints such as LogD, this broad coverage may be sufficient, leaving little systematic gain for uncertainty ranking. Random sampling is therefore not a trivial baseline in clustered molecular pools: it provides broad stochastic chemical diversity. The observed diversity-aware point estimates may be related to their broader batch coverage, but this post-hoc association is not causal evidence.

The method-level diagnostics in Table 5 sharpen this interpretation. The uncertainty–error correlations are similar across acquisition functions, but the outcomes are not. Noise-Aware acquisition has the highest median uncertainty–error correlation (0.081) yet the worst median final ΔRMSE (+0.056), while Core-set has slightly lower uncertainty quality (0.077) but the best median next-round outcome (−0.001, the only negative value in that column) and a final outcome (−0.005) tied with COVDROP at the reported precision. BADGE and Core-set also lose less cluster coverage relative to random than the pointwise uncertainty and covariance-ranking methods. The practical hypothesis is that ranking uncertain molecules is not enough; a successful acquisition function would also need to assemble a batch that expands useful chemical coverage. The present diagnostics do not establish that doing so causes improved generalization.

Validation-only conformal calibration is used here to construct empirically calibrated intervals and report observed test coverage, rather than as an acquisition signal. Because validation and test occupy different held-out chemotype clusters, standard exchangeability guarantees need not apply. An empirically calibrated interval width is a natural future acquisition criterion, but using it in the present benchmark would change the locked acquisition protocol and would require a separate, prospective comparison. The current results therefore use empirically calibrated intervals as an evaluation safeguard, not as evidence that interval width itself improves selection.

Caco-2 Efflux Ratio requires an additional calibration caveat because its Box–Cox inverse transform can amplify upper-tail interval bounds in original units. Mean observed test coverage of the empirically calibrated intervals remains close to the target for both backends (PICP ≈ 0.975), but roughly one quarter of Efflux prediction intervals reach the 10^6^ upper cap. This cap is a numerical safeguard applied when inverting the Box–Cox transform, not a property of the data: original-unit outputs are clipped to |*y |≤* 10^6^ to keep the inverse transform finite. It is therefore about 10^4^ times the largest observed efflux ratio (105.64, Table 1), and an interval bound can reach it because the Box–Cox inverse transform can strongly amplify upper-tail bounds. Efflux uses the *c*–*τ* calibration described in Methods; its interval bounds are not constrained by the observed maximum. Reaching the cap indicates an uninformatively wide interval, not a predicted efflux ratio of 10^6^. We therefore interpret interval-width results for this endpoint only alongside coverage, and use RMSE/MAE-based paired comparisons for the main active-learning claims.

The endpoint-level diagnostics provide descriptive hypotheses rather than additional inferential claims. KSOL has aggregate uncertainty–error correlation 0.175 and median next-round ΔRMSE 0.120; Caco-2 Papp A→B has the highest correlation (0.214). LogD and HLM CLint have weaker ranking (0.036 and 0.031) and reduced cluster coverage. MLM CLint is volatile, Efflux has modest ranking (0.095), and MPPB and MBPB remain mixed. These post-hoc patterns may guide endpoint-specific replication and the design of diversity-aware acquisition studies.

### Endpoint-Level Interpretation

Table 6 summarizes descriptive endpoint-level patterns under the strict chemotype cluster split. KSOL has the broadest favorable point estimates, random remains competitive on LogD and HLM CLint, and the remaining endpoints show method-specific effects. Together, these patterns guide targeted follow-up across the full panel.

**Table 6:** Endpoint-level interpretation of final active-learning behavior.

| Endpoint | Empirical pattern | Interpretation |
| --- | --- | --- |
| LogD | Mostly random-competitive; effects are predominantly positive (worse). | The initial/random batches may cover the relevant chemistry well; uncertainty ranking adds little descriptively. |
| KSOL | Broadest negative ΔRMSE point estimates. | A priority for replicating the favorable acquisition pattern. |
| HLM CLint | Positive effects across methods and backends. | Random provides a competitive reference; Core-set is closest in mean RMSE. |
| MLM CLint | Large effects in both directions. | Core-set is favorable under both backends; seed variability warrants replication. |
| Caco-2 Papp A→B | Several negative covariance-aware point estimates. | Batch complementarity is a hypothesis for targeted replication. |
| Caco-2 Efflux Ratio | Mixed, method-dependent effects. | Coverage remains near target, but transformed-tail interval width requires caution. |
| MPPB | Several negative point estimates under both backends. | Effects depend on the backend and method; replication should retain this pairing. |
| MBPB | Mixed, mostly small effects; random remains competitive. | Interpret paired effects with the documented round 9–10 aggregation. |

### Practical Implications

For ADMET campaigns expected to face similarly strict chemotype shift, the results suggest a conservative workflow. Random acquisition should remain the mandatory baseline because it provides broad chemical-space coverage and is difficult to improve consistently. An active-learning method should be adopted only after an endpoint-specific pilot demonstrates a reproducible gain against that baseline, using the same assay budget and split structure as the intended campaign. Diversity-aware criteria remain candidates for endpoint-specific pilots because they explicitly target the batch concentration observed in the post-hoc diagnostics, although this benchmark does not establish that broader coverage causes better outcomes. Deep Ensembles provide stronger matched predictive accuracy at exactly five times the trained-model count, but wall-clock cost does not follow that ratio once stochastic inference is included. The cheaper MC Dropout acquisition functions run faster than any Deep Ensemble configuration, while the most expensive run slower, so compute should be budgeted per acquisition function rather than per backend. Prediction intervals should be judged jointly by coverage and width, especially for transformed-tail endpoints such as Efflux.

These implications are deliberately narrower than a claim about all ADMET active learning: they apply to the strict chemotype cluster split, where evaluation chemotypes are unavailable for acquisition. In practice, the acquisition rule should be prespecified, random matched controls should be retained, calibration should use validation data only, and test-set diagnostics should be treated as post-hoc explanation rather than as a tuning signal.

### Limitations

Several limitations define the scope of these conclusions. The active-learning benchmark uses one molecular architecture, one curated dataset, and a fixed batch size of 48; results may change with alternative representations, datasets, assay budgets, or batch sizes. The fixed-data Random Forest check broadens predictor comparison but does not provide Random-Forest uncertainty or acquisition trajectories. The active-learning benchmark’s initialization is target-informed: the 200 starting molecules are selected from the train pool using their target labels through 12-bin regression stratification. Although no validation or test labels are used, this reduces target-range imbalance and is not directly realizable in a prospective unlabeled campaign. The resulting starting set may be unusually informative and may help explain why random acquisition is competitive; uniform-random and label-free diversity initializations require a separate ablation. More generally, the retrospective pool-based design cannot reproduce all operational constraints and distributional changes of a prospective wet-lab campaign, and industrial experience indicates that retrospective model rankings do not always persist prospectively ^6^. In addition, MBPB lacks matched round-10 artifacts for seeds 0, 5, and 6 (three of ten seeds); analyses use the final common available round for those matched cases, which weakens the ideal fixed-round comparison for this endpoint. MGMB is retained for protocol and split audits but excluded from the paper-level claims because its split is too small. Finally, the Caco-2 Efflux Ratio analysis now uses the same 200-molecule initial labeled set as the other paper endpoints; earlier 100-molecule outputs were replaced by the validated 200-molecule rerun, so the initial-set asymmetry is not part of the reported comparison. These limitations motivate validation across architectures, datasets, budgets, and prospective experiments.

The single-architecture active-learning design and single dataset also limit external validity: the endpoint-dependent pattern may reflect the representation, assay practices, chemical composition of OpenADMET ExpansionRx, and the deliberately strict cluster shift. Settings in which the acquisition pool overlaps more closely with deployment chemistry may yield different conclusions. Because endpoints are modeled independently, the study does not test whether multi-task sharing would change sample efficiency or acquisition priorities. The fixed batch size of 48 and retrospective pool do not represent all plate sizes, assay costs, synthesis constraints, failed measurements, or changing candidate pools encountered prospectively. These constraints motivate cost-aware and prospective validation rather than direct deployment of one acquisition rule.

The uncertainty–actionability analysis is descriptive and post-hoc, so its proposed associations and hypotheses—likelihood misspecification, warm-start effects, representation limitations, and error associated with chemotype shift—are not causal findings. Similarly, the NN-Cov results apply only to the disclosed noise-normalized covariance method. Finally, the missing MBPB round-10 outputs and the small audit-only

MGMB split limit those endpoint-specific conclusions; Efflux interval widths also remain sensitive to the Box–Cox inverse transform and must be interpreted alongside coverage.

Statistical resolution is limited by ten paired seeds and the breadth of the comparison families. No acquisition-versus-random RMSE comparison meets the Benjamini–Hochberg threshold of *q <* 0.05 across 48 tests per backend. Adjustment across this many comparisons can reduce sensitivity to modest effects; the absence of a discovery establishes neither equivalence nor an acquisition benefit. Effect sizes and seed-level uncertainty therefore guide targeted replication, with full raw and adjusted results retained in the Supplementary Materials.

### Future Work

The most direct methodological extension is to use empirically calibrated interval width as an acquisition signal. This should be evaluated as a separate nested protocol: calibration must use validation data only, interval width must be computed before querying, and the comparison must retain the same matched random baseline. Such an experiment would directly test whether empirical calibration can turn the uncertainty signal into more actionable selection.

Several controlled ablations could test which design choices alter the observed behavior. Warm-starting versus cold-starting could test whether repeated fine-tuning narrows model diversity; reducing the number of MC passes could quantify the accuracy–runtime trade-off; and batch-size experiments could establish whether the advantage of random or diversity-aware selection changes at realistic plate sizes. Although the fixed-data Random Forest check compares predictors after a common random trajectory, full representation ablations should compare acquisition trajectories driven by the current GNN, classical ensembles, a pure GNN, and pretrained molecular representations.

The benchmark also motivates multi-task and cost-aware active learning. A multi-task model could share information across clearance, permeability, and binding endpoints while selecting molecules for a complete ADMET profile. A cost-aware objective could account for assay time, monetary expense, failed measurements, or information per label rather than treating every endpoint and molecule as equally costly. Hybrid criteria combining uncertainty with Core-set or COVDROP-style diversity may be useful because the present results suggest that uncertainty quality and batch coverage are complementary.

Finally, prospective wet-lab validation is needed to establish operational value. KSOL is a reasonable first target because it shows the broadest favorable retrospective point estimates, while Efflux and MBPB would test whether the identified failure modes persist in difficult or data-limited settings. Future studies should vary the degree of scaffold shift, extend the framework to classification endpoints such as hERG or Ames, and develop a formal theory of the uncertainty–actionability gap.

## Conclusion

This study provides a protocol-matched framework for comparing active-learning strategies and uncertainty backends across eight ADMET endpoints. Under the strict chemotype cluster split, acquisition effects are endpoint- and method-dependent: solubility shows the broadest favorable point estimates, selected diversity-aware strategies are promising candidates for replication, and random acquisition remains a competitive reference across the panel. Deep Ensembles have lower mean RMSE in most matched backend comparisons, with supported differences concentrated in particular endpoints. The fixed-data Random Forest comparison further highlights complementary strengths across predictors and error metrics.

The combined findings motivate evaluating uncertainty ranking, batch diversity, prediction accuracy, and computational cost together. They support endpoint-specific pilots with matched random controls and validation-only calibration, followed by prospective evaluation in chemistry representative of the intended campaign. Prediction-interval width should be interpreted alongside empirical coverage, particularly for skewed transformed endpoints such as Caco-2 Efflux Ratio. The framework and paired analyses provide a reproducible basis for these comparisons and for developing acquisition strategies suited to different ADMET tasks.

## Supporting Information

The accompanying Supplementary Materials report the locked protocol table, endpoint inclusion audit, compact split and output completeness checks, representative learning curves, compact paired seed-level inference summaries, the fixed-data Random Forest comparison, uncertainty-actionability diagnostics, compute-aware summaries, and validation status. Exhaustive per-cell records are retained as machine-readable analysis artifacts.

## Data and Code Availability

Code for the active-learning experiments and analysis is available at https://github.com/VikiShmiki/Active-Learning-ADMET. The OpenADMET ExpansionRx data are publicly available from the OpenADMET Hugging Face collection at https://huggingface.co/openadmet. Large generated outputs and model artifacts are not versioned in the repository.

## Abbreviations

pADMET: absorption, distribution, metabolism, excretion, and toxicity
AL: active learning
GNN: graph neural network
MC: Monte Carlo
RMSE: root mean squared error
MAE: mean absolute error
PICP: prediction interval coverage probability
NLL: negative log-likelihood

## Notes

https://github.com/VikiShmiki/Active-Learning-ADMET

### Competing Interest Statement

The authors have declared no competing interest.

