## Supplementary Materials for "When Does Active Learning Improve ADMET Prediction? A Multi-Endpoint Benchmark"

#### Overview

This concise supplement documents a controlled stress test under strict chemotype-cluster extrapolation. Whole held-out test chemotypes are absent from the acquisition pool, so no method can query molecules from the chemistry on which it is evaluated; this is the difficult end of the distribution-shift spectrum, not a universal model of every ADMET campaign. It contains the protocol, split and completeness audit, compact statistical summaries for the four lower-is-better metrics cited in the manuscript in both the versus-random and cross-backend directions, representative learning curves, a fixed-data Random Forest predictor comparison, backend-split uncertainty–actionability diagnostics, and an annotated aggregate compute report for the eight-endpoint, protocol-matched benchmark. Exhaustive per-seed and per-cell tables remain available as machine-readable analysis artifacts in the accompanying repository output and are not reproduced here to keep the print supplement focused.

All results use the final validated paper-ready roots. No training, acquisition, split, calibration, or result-filtering inputs were changed to prepare these materials. Test data were never used for acquisition, calibration, tuning, early stopping, model selection, or run filtering. Reported test intervals are described as empirically calibrated intervals: formal conformal coverage guarantees require exchangeability, whereas validation and test occupy different held-out chemotype clusters here, so that assumption need not hold. Coverage is therefore evaluated empirically under the benchmark's chemotype shift.

#### Locked Protocol

**Table S1:** Locked benchmark protocol. The Efflux Ratio endpoint uses the validated 200-molecule rerun.

| Item | Value | Note |
|---|---|---|
| Dataset | OpenADMET ExpansionRx | Independent regression per endpoint |
| Paper endpoints | Eight validated endpoints | MGMB retained for audit only |
| Split mode | Cluster | Shared immutable splits |
| Seeds | 0–9 | Matched across backends |
| Acquisition methods | Random plus six non-random methods | Batch size 48 |
| Initial labeled size | 200 molecules | Target-informed 12-bin stratification within train pool |
| Maximum rounds | 10 | MBPB seeds 0, 5, and 6 stop at round 9 |
| Target coverage | 0.95 | Validation-only conformal calibration |
| Conformal bins | 4; minimum size 12 | Recomputed each round |
| Clustering | MiniBatchKMeans, $k = \lfloor\sqrt{n}\rfloor$ clipped to $[10, 200]$ | $n$ is the endpoint-specific featurizable unique-SMILES population (Table S3) |
| MC Dropout | $T_{\text{screen}} = 8$ screening; $T_{\text{acq}} = 96$ acquisition; $T_{\text{val}} = T_{\text{test}} = 128$ evaluation passes | Dropout active at inference |
| Deep Ensemble | 5 deterministic members | Dropout disabled at inference |
| Shortlist multiplier | 10 | Expensive acquisition scoring |

Table S2 lists the per-endpoint settings. The initial labeled size is 200 molecules for every paper endpoint, including Caco-2 Efflux Ratio, whose earlier 100-molecule outputs were superseded by the validated 200-molecule rerun. Within each training pool, these molecules are selected using the train-pool target labels through seeded stratification over 12 quantile-style regression bins. This target-informed retrospective construction reduces target-range imbalance; validation and test labels are never used. It is not directly realizable in a prospective campaign with a genuinely unlabeled pool, may provide an unusually informative starting set, and may help explain why subsequent random acquisition is competitive. Uniform-random and label-free ECFP diversity initializations are therefore reserved for the planned initialization ablation. MGMB is retained only for split and protocol audits; its split is too small to support paper-level claims and it uses a 64-molecule initial set.

**Table S2:** Endpoint settings and inclusion status. Initial $n$ is the *effective* initial labeled-set size realized in the locked split artifacts, and equals the requested 200 molecules for all eight paper endpoints. MGMB requests 200 but resolves to 64 under the reserved-pool rule for very small splits, which is one reason it is excluded from paper-level claims.

| Endpoint | Aliases | Units | Initial $n$ | Target transform | Calibration | Status |
|---|---|---|---|---|---|---|
| LogD | — | — | 200 | identity | picp_original_mondrian | included |
| KSOL | — | $\mu$M | 200 | identity | picp_original_mondrian | included |
| HLM CLint | — | mL/min/kg | 200 | yeo_johnson | picp_original_mondrian | included |
| MLM CLint | — | mL/min/kg | 200 | yeo_johnson | picp_original_mondrian | included |
| Caco-2 Papp A→B | Caco-2 Papp A>B; Caco-2 Permeability Papp A>B | $10^{-6}$ cm/s | 200 | identity | picp_original_mondrian | included |
| Caco-2 Efflux Ratio | Caco-2 Permeability Efflux; Caco-2 Permeability Efflux Ratio | — | 200 | boxcox_shift | ctau_mondrian | included |
| MPPB | — | % unbound | 200 | yeo_johnson | picp_original_mondrian | included |
| MBPB | — | % unbound | 200 | yeo_johnson | picp_original_mondrian | included |
| MGMB | — | % unbound | 64 | yeo_johnson | picp_original_mondrian | excluded (audit only) |

### Splits and Completeness

Clustering and splitting were performed independently for each endpoint. Whole clusters are assigned to exactly one partition, so test chemotypes are absent from both the training and acquisition pools. This stringent separation makes the benchmark a controlled extrapolation stress test and limits its conclusions to that regime. The split audit below reports mean (range) across the ten matched seeds; full per-seed records are retained in the accompanying CSV artifact. Cluster counts are fixed across seeds for a given endpoint, while partition sizes vary because whole clusters are assigned to partitions.

Table S3 reports the deduplicated, featurizable population $n$ that clustering actually operates on, so that the locked rule $k = \lfloor\sqrt{n}\rfloor$ (clipped to $[10, 200]$) is directly checkable per endpoint. This population is smaller than the raw labeled-row count in Table 1 of the manuscript because the dataset is de-duplicated on SMILES before splitting and because molecules whose fingerprints or descriptors cannot be computed are dropped. It is constant across seeds — only the partition assignment changes with the seed — and it is exactly the sum of the train-pool, validation, and test partitions. Reported cluster counts equal $\lfloor\sqrt{n}\rfloor$ for all eight endpoints with no exceptions.

**Table S3:** Compact split audit. Partition sizes are mean (minimum–maximum) across the ten seeds. Unique $n$ is the deduplicated featurizable population used for clustering; it is seed-invariant, and $\lfloor\sqrt{n}\rfloor$ reproduces the reported cluster count for every endpoint.

| Endpoint | Unique $n$ | Train pool | Validation | Test | Initial | Clusters | $\lfloor\sqrt{n}\rfloor$ |
|---|---|---|---|---|---|---|---|
| LogD | 5039 | 3349 (3276–3406) | 637 (606–690) | 1054 (1022–1125) | 200 | 70 | 70 |
| KSOL | 5128 | 3386 (3238–3451) | 664 (640–748) | 1078 (1030–1217) | 200 | 71 | 71 |
| HLM CLint | 3759 | 2478 (2371–2516) | 494 (462–577) | 787 (761–888) | 200 | 61 | 61 |
| MLM CLint | 4522 | 2971 (2819–3047) | 577 (543–657) | 974 (906–1056) | 200 | 67 | 67 |
| Caco-2 Papp A→B | 2157 | 1416 (1380–1441) | 284 (271–296) | 457 (435–506) | 200 | 46 | 46 |
| Caco-2 Efflux Ratio | 2161 | 1412 (1335–1457) | 289 (262–331) | 460 (442–512) | 200 | 46 | 46 |
| MPPB | 1302 | 848 (811–874) | 172 (160–180) | 283 (261–315) | 200 | 36 | 36 |
| MBPB | 975 | 632 (600–647) | 131 (122–142) | 212 (196–242) | 200 | 31 | 31 |

**Table S4:** Completeness audit for final matched outputs.

| Backend | Output kind | Observed | Expected | Missing |
|---|---|---|---|---|
| Deep Ensemble | initial acquisition | 560 | 560 | 0 |
| Deep Ensemble | metadata | 6139 | 6160 | 21 |
| Deep Ensemble | prediction | 6139 | 6160 | 21 |
| Deep Ensemble | selected acquisition | 5579 | 5600 | 21 |
| MC Dropout | initial acquisition | 560 | 560 | 0 |
| MC Dropout | metadata | 6139 | 6160 | 21 |
| MC Dropout | prediction | 6139 | 6160 | 21 |
| MC Dropout | selected acquisition | 5579 | 5600 | 21 |

The only missing matched outputs are MBPB round 10 for seeds 0, 5, and 6 across all seven methods and both backends. Paired analyses therefore use the final common round for those matched cases. This limitation is disclosed in the manuscript.

**Table S5:** Matched missing outputs. Missing records are paired across backends.

| Backend | Kind | Endpoint | Round | Seeds | Methods | Missing |
|---|---|---|---|---|---|---|
| Deep Ensemble | metadata | MBPB | 10 | 0, 5, 6 | all 7 methods | 21 |
| Deep Ensemble | prediction | MBPB | 10 | 0, 5, 6 | all 7 methods | 21 |
| Deep Ensemble | selected acquisition | MBPB | 10 | 0, 5, 6 | all 7 methods | 21 |
| MC Dropout | metadata | MBPB | 10 | 0, 5, 6 | all 7 methods | 21 |
| MC Dropout | prediction | MBPB | 10 | 0, 5, 6 | all 7 methods | 21 |
| MC Dropout | selected acquisition | MBPB | 10 | 0, 5, 6 | all 7 methods | 21 |

### Representative Learning Curves

Rather than reproduce the full set of near-duplicate metric panels, Figures S1 and S2 give the complete RMSE learning curves for all eight paper endpoints under both backends, split across two pages for legibility. The corresponding MAE, PICP95, ACE95, and interval-width curves are retained in the machine-readable supplementary artifact set; Efflux interval widths should always be interpreted together with coverage because the Box–Cox inverse transform can amplify upper-tail bounds.

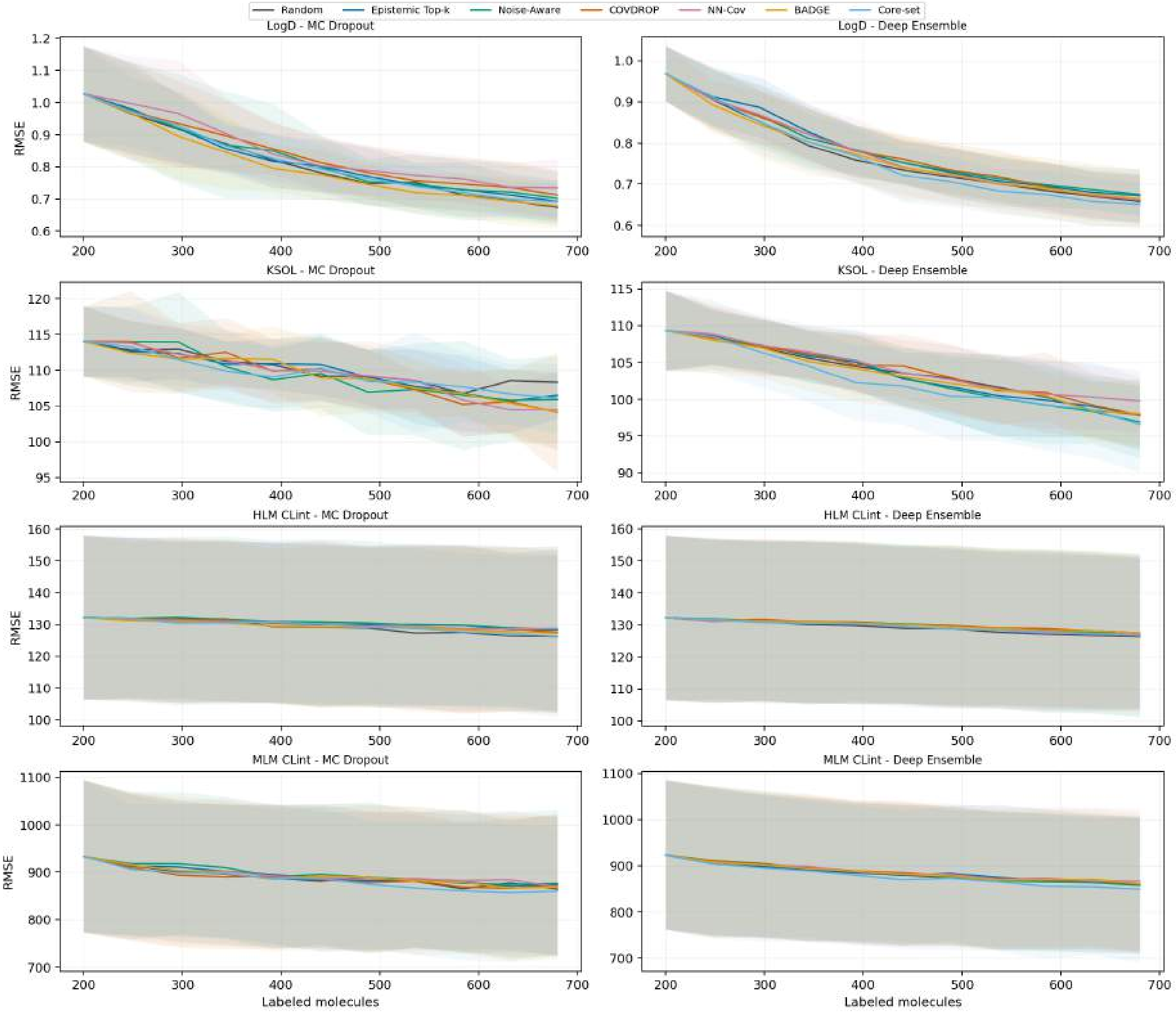


**Figure S1:** RMSE learning curves, part 1 of 2: LogD, KSOL, HLM CLint, and MLM CLint. Columns are the two uncertainty backends. Lines show seed means; shaded bands show one standard deviation for non-random methods.

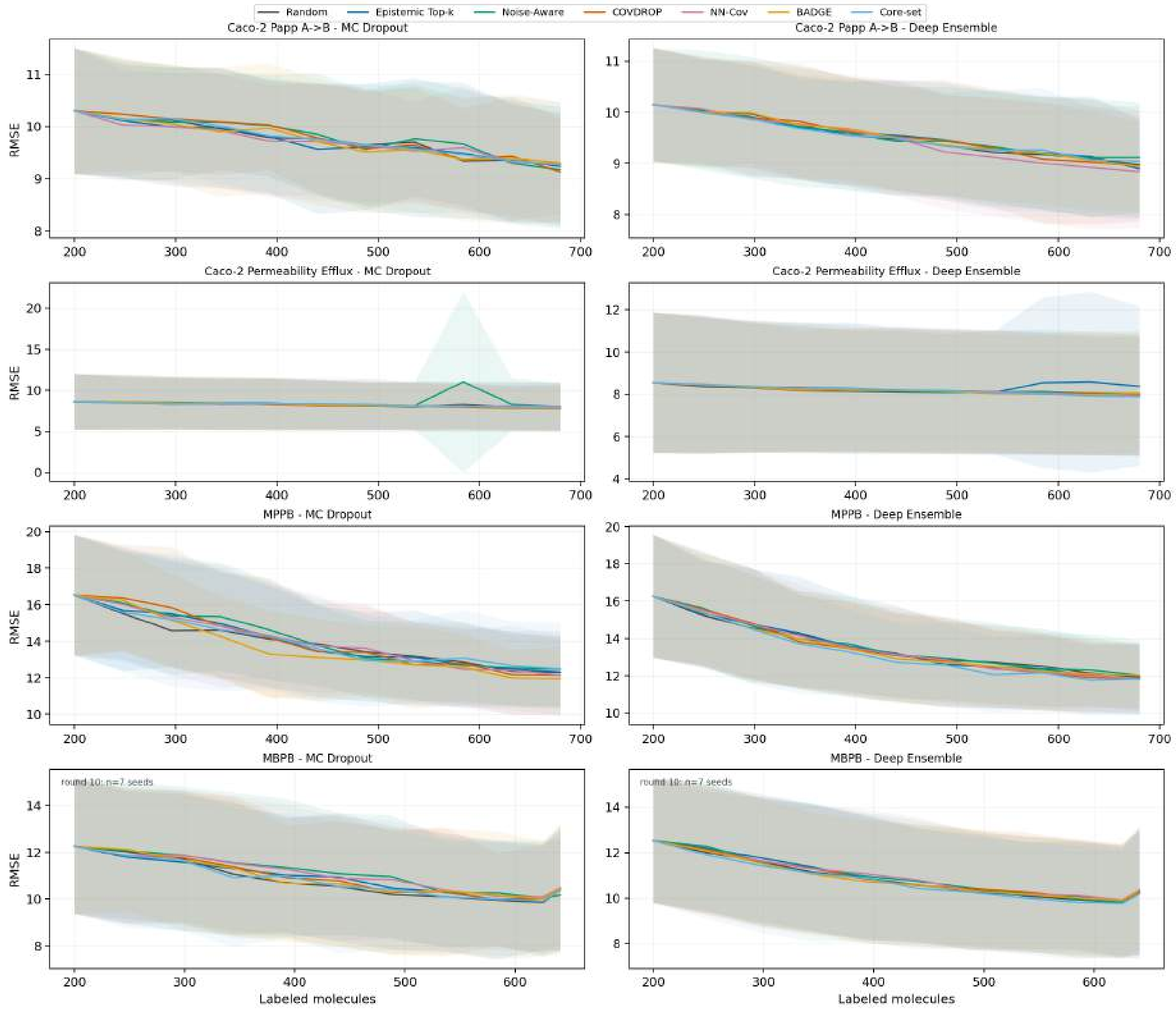


**Figure S2:** RMSE learning curves, part 2 of 2: Caco-2 Papp A→B, Caco-2 Efflux Ratio, MPPB, and MBPB. Columns are the two uncertainty backends. Lines show seed means; shaded bands show one standard deviation for non-random methods. MBPB has seven seeds at round 10 because seeds 0, 5, and 6 stop at round 9.

### Paired Seed-Level Inference

The mean of the ten paired seed-level metric differences is the primary estimand. Bootstrap resampling estimates uncertainty over ten paired seed-level differences; $B$ = 10,000 controls Monte Carlo precision and does not increase the effective number of experimental replicates. Raw p-values are exact two-sided paired sign-flip tests. Benjamini–Hochberg (BH) correction is applied to prespecified families: 48 acquisition-versus-random RMSE tests separately per backend, a pooled 96-test RMSE sensitivity family, 48 tests separately per backend for each secondary metric, and 56 cross-backend tests per metric. Exact sign tests are retained as a sensitivity analysis in the machine-readable aggregate files. MBPB uses the final common round: seven seeds contribute round 10 and seeds 0, 5, and 6 contribute round 9.

**Table S6:** Primary within-backend final-RMSE results. Effect directions count negative (better than random) and positive (worse than random) mean paired-seed deltas descriptively. Raw and BH columns count exact sign-flip $p < 0.05$ and within-family $q < 0.05$ decisions.

| Backend | Family size | Negative | Positive | Raw rejects | BH rejects |
|---|---|---|---|---|---|
| MC Dropout | 48 | 14 | 34 | 5 | 0 |
| Deep Ensemble | 48 | 19 | 29 | 2 | 0 |
| Pooled sensitivity | 96 | 33 | 63 | 7 | 0 |

The seven raw within-backend RMSE rejections all have positive deltas. Under MC Dropout these are HLM CLint NN-Cov ($\Delta = +2.619$, 95% seed CI $[+1.438, +3.900]$, $p = 0.0039$, $q = 0.0938$), HLM CLint Epistemic ($+2.037$ $[+1.074, +3.082]$, $p = 0.0039$, $q = 0.0938$), HLM CLint Noise-Aware ($+2.211$ $[+0.772, +3.891]$, $p = 0.0176$, $q = 0.2109$), LogD NN-Cov ($+0.061$ $[+0.029, +0.090]$, $p = 0.0098$, $q = 0.1563$), and LogD COVDROP ($+0.039$ $[+0.013, +0.064]$, $p = 0.0234$, $q = 0.2250$). Under Deep Ensemble they are LogD Noise-Aware ($+0.016$ $[+0.006, +0.026]$, $p = 0.0234$, $q = 0.5625$) and MBPB NN-Cov ($+0.128$ $[+0.047, +0.214]$, $p = 0.0176$, $q = 0.5625$). Thus, no acquisition-versus-random RMSE result remains significant after adjustment. The secondary within-backend MAE, NLL, ACE95, PICP95, and interval-width families likewise have zero BH discoveries. PICP95 is interpreted by distance from 0.95 through ACE95, not as larger-is-better; interval-width inference is conditional on adequate coverage.

**Table S7:** Cross-backend final-round results by metric. Negative effects favor Deep Ensemble for the lower-is-better metrics. Raw and adjusted columns count exact sign-flip and within-metric BH decisions. Interval-width results are conditional on adequate empirical coverage.

| Metric | Family size | Negative | Positive | Raw rejects | BH rejects |
|---|---|---|---|---|---|
| RMSE | 56 | 45 | 11 | 15 | 7 |
| MAE | 56 | 52 | 4 | 15 | 5 |
| NLL | 56 | 54 | 2 | 38 | 29 |
| ACE95 | 56 | 28 | 28 | 1 | 0 |
| PICP95 | 56 | 25 | 31 | 3 | 0 |
| AvgIntWidth | 56 | 46 | 10 | 9 | 3 |

For RMSE, the seven BH discoveries all favor Deep Ensemble: KSOL under Random, Epistemic, Noise-Aware, COVDROP, BADGE, and Core-set, plus NN-Cov on Caco-2 Papp A→B. KSOL effects range from $-6.085$ to $-10.397$; their 95% seed-bootstrap intervals exclude zero and their BH q-values are 0.0182, except COVDROP ($q = 0.0469$). Papp NN-Cov has $\Delta = -0.447$, 95% CI $[-0.770, -0.208]$, exact $p = 0.0020$, and $q = 0.0182$. These cross-backend findings concern predictor performance; they do not imply that non-random acquisition beats random.

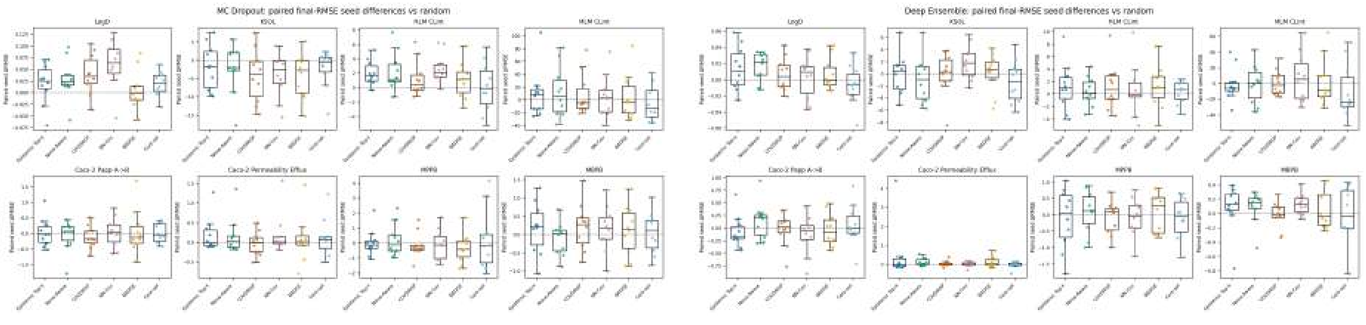


**(a)** MC Dropout.

**(b)** Deep Ensemble.

**Figure S3:** Paired final-RMSE seed differences for every acquisition-versus-random comparison. Points are the ten experimental seeds; boxes summarize those seed-level differences. Negative values favor acquisition. These distributions, rather than the number of test molecules, determine inferential uncertainty.

Full per-cell records report the effect size, seed-bootstrap interval, exact sign-flip p-value, BH q-value, raw and adjusted decisions, exact sign-test sensitivity result, seed identifiers and differences, family size, $B$, and actual round range. They are regenerated in the accompanying machine-readable supplementary artifact set.

#### Fixed-Data Random Forest Baseline

For each endpoint and seed, a classical Random Forest was fit to the exact final labeled set reconstructed from the random-acquisition trajectory. It uses a chiral 2,048-bit ECFP4 fingerprint (radius 2), nine RDKit descriptors, training-only median imputation, and prespecified hyperparameters (500 trees, bootstrap sampling, square-root feature sampling, and minimum leaf size one). Validation metrics are diagnostic only; no validation- or test-driven tuning is performed. Test molecules and labels are aligned one-to-one with each GNN random-baseline artifact. This comparison evaluates predictor competitiveness at fixed acquired data and must not be interpreted as a Random-Forest uncertainty-backend or active-learning comparison.

The effect is Random Forest minus GNN, so negative RMSE or MAE favors the Random Forest. In the prespecified 16-test family per metric, adjusted RMSE results favor the Random Forest in 5 comparisons and a GNN in 6, with 5 null; adjusted MAE favors the Random Forest in 1 and a GNN in 13, with 2 null. Table S8 gives all endpoint-level effects and q-values. The accompanying CSVs retain all 80 endpoint–seed fits, 320 paired metric rows, seed-bootstrap intervals, exact sign-flip p-values, and multiplicity-adjusted endpoint summaries.

**Table S8:** Fixed-data Random Forest minus GNN effects on the matched final random-trajectory labeled sets. Negative $\Delta$ favors the Random Forest; positive $\Delta$ favors the indicated GNN. Reported q-values use BH correction within the prespecified 16-test metric family; exact paired sign-flip raw p-values are retained in the accompanying CSV.

| | MC Dropout GNN | | | | Deep Ensemble GNN | | | |
|---|---|---|---|---|---|---|---|---|
| Endpoint | $\Delta$RMSE | q | $\Delta$MAE | q | $\Delta$RMSE | q | $\Delta$MAE | q |
| LogD | +0.217 | 0.0045 | +0.188 | 0.0026 | +0.231 | 0.0045 | +0.202 | 0.0026 |
| KSOL | -8.779 | 0.0045 | -2.188 | 0.4258 | +1.619 | 0.1853 | +4.894 | 0.0168 |
| HLM CLint | -4.588 | 0.0045 | +7.428 | 0.0026 | -4.720 | 0.0104 | +7.603 | 0.0026 |
| MLM CLint | -20.283 | 0.0573 | +70.904 | 0.0026 | -15.374 | 0.1010 | +78.296 | 0.0026 |
| Caco-2 Papp A→B | -0.648 | 0.0045 | -0.251 | 0.0446 | -0.381 | 0.0078 | -0.105 | 0.3229 |
| Caco-2 Efflux Ratio | +0.075 | 0.7229 | +0.834 | 0.0026 | +0.019 | 0.8965 | +0.834 | 0.0026 |
| MPPB | +1.695 | 0.0114 | +2.112 | 0.0026 | +2.043 | 0.0045 | +2.336 | 0.0026 |
| MBPB | +0.715 | 0.0114 | +1.287 | 0.0026 | +0.824 | 0.0045 | +1.338 | 0.0026 |

The mixed RMSE pattern and the stronger GNN MAE pattern do not support a universal architecture ranking. They also do not alter the active-learning comparison, because every Random Forest uses data selected by the existing random trajectory rather than selecting its own molecules.

#### Uncertainty and Actionability Diagnostics

The actionability analysis is post-hoc and descriptive. It uses test-set errors only after all active-learning runs were complete. Random batches provide the empirical diversity baseline because uncertainty for all unselected molecules was not stored.

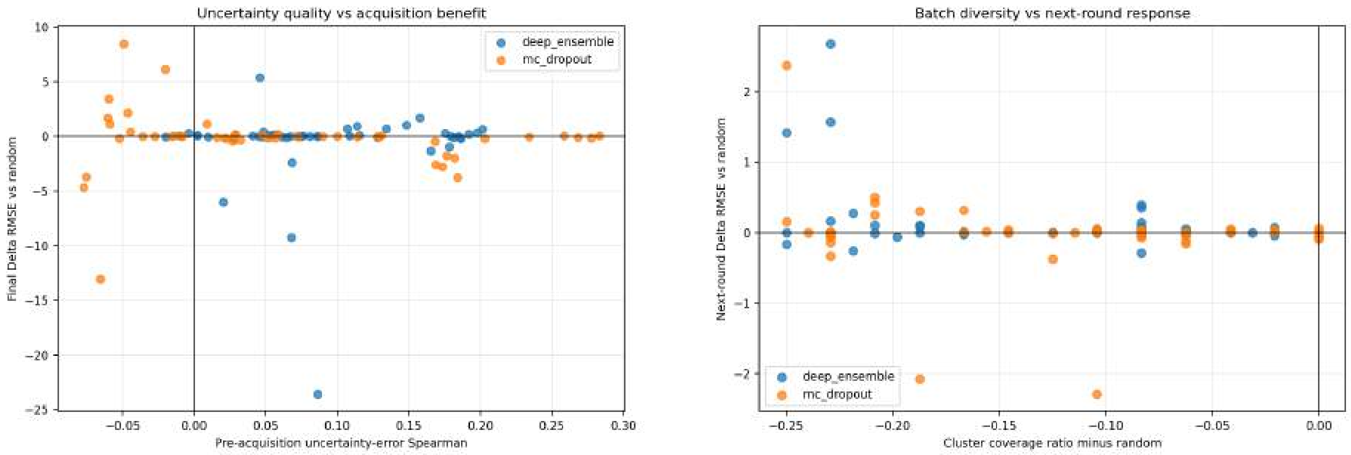


**(a)** Uncertainty quality versus final gain.

**(b)** Batch diversity versus next-round response.

**Figure S4:** Representative uncertainty–actionability diagnostics. Negative RMSE deltas favor active learning over matched random acquisition.

Table S9 is the backend-split companion to Table 5 of the manuscript, which pools the two backends. It uses identical column definitions and the identical set of matched cells, split by backend, so that the backend-specific figures quoted in the Discussion are directly checkable. Deep Ensemble ranks high-error molecules better than MC Dropout on every uncertainty-quality column, yet its median acquisition outcome versus random is still slightly positive; this post-hoc association is consistent with, but does not causally establish, an uncertainty–actionability gap. Batch-composition losses are backend-invariant: both backends give up the same median cluster coverage (−0.125, about six of 48 molecules) and show the same tail enrichment (+0.021). The one column that differs sharply is aleatoric-fraction enrichment, which reflects the different epistemic-to-aleatoric variance scales of the two backends rather than a difference in what the acquisition functions select; it should not be compared across backends.

**Table S9:** Backend-split post-hoc actionability diagnostics: the backend-level companion to manuscript Table 5. Medians over matched endpoint, seed, and round cells for the six non-random acquisition methods (4,782 cells per backend, 9,564 pooled). $\rho_{\mathrm{epi},|e|}$ is the pre-acquisition Spearman correlation between epistemic uncertainty and absolute test error; Enrich. is the top-decile error-enrichment ratio; Overlap is the top-uncertainty/top-error decile overlap (0.10 expected by chance). RMSE and batch-composition deltas are relative to the matched random batch; negative RMSE deltas are better than random.

| Backend | Method | $\rho_{\mathrm{epi},\lvert e\rvert}$ | Enrich. | Overlap | Next ΔRMSE | Final ΔRMSE | ΔCluster cov. | ΔTail | ΔAleatoric |
|---|---|---|---|---|---|---|---|---|---|
| Deep Ensemble | Epistemic Top-$k$ | 0.087 | 1.164 | 0.123 | +0.0007 | +0.006 | −0.167 | +0.021 | +0.803 |
| | Noise-Aware | 0.092 | 1.155 | 0.117 | +0.0031 | +0.035 | −0.167 | +0.000 | +0.795 |
| | COVDROP | 0.092 | 1.144 | 0.115 | +0.0052 | +0.013 | −0.167 | +0.021 | +0.805 |
| | NN-Cov | 0.092 | 1.139 | 0.115 | +0.0062 | +0.017 | −0.146 | +0.021 | +0.806 |
| | BADGE | 0.091 | 1.153 | 0.111 | +0.0039 | +0.038 | −0.062 | +0.021 | +0.846 |
| | Core-set | 0.097 | 1.158 | 0.115 | +0.0002 | −0.011 | −0.062 | +0.000 | +0.899 |
| | **All six pooled** | **0.092** | **1.154** | **0.115** | **+0.0033** | **+0.017** | **−0.125** | **+0.021** | **+0.824** |
| MC Dropout | Epistemic Top-$k$ | 0.051 | 1.111 | 0.109 | +0.0033 | +0.033 | −0.167 | +0.021 | −0.030 |
| | Noise-Aware | 0.068 | 1.134 | 0.111 | +0.0010 | +0.056 | −0.167 | +0.000 | −0.031 |
| | COVDROP | 0.056 | 1.110 | 0.109 | +0.0000 | −0.081 | −0.146 | +0.021 | −0.025 |
| | NN-Cov | 0.064 | 1.136 | 0.111 | +0.0140 | +0.072 | −0.167 | +0.000 | −0.042 |
| | BADGE | 0.059 | 1.117 | 0.109 | +0.0039 | −0.031 | −0.021 | +0.021 | −0.035 |
| | Core-set | 0.055 | 1.098 | 0.109 | −0.0038 | +0.001 | −0.062 | +0.021 | −0.054 |
| | **All six pooled** | **0.059** | **1.118** | **0.109** | **+0.0035** | **+0.021** | **−0.125** | **+0.021** | **−0.037** |

Endpoint-level actionability summaries and the remaining method-by-endpoint diagnostics are retained in the machine-readable artifact set; the main manuscript reports the aggregate patterns needed for interpretation.

### Compute-Aware Reporting

The comparison is protocol-matched, not compute-matched. MC Dropout trains one stochastic model per method, seed, endpoint, and round; Deep Ensemble trains five deterministic members. Deep Ensembles therefore train exactly five times as many models, but wall-clock cost does not follow that ratio, because MC Dropout performs many more stochastic inference passes per trained model. The MC Dropout range straddles the Deep Ensemble range rather than sitting above or below it: MC Dropout spans 10.10–14.71 h across the seven acquisition methods against Deep

Ensemble's much narrower 11.19–12.22 h. The cheapest MC Dropout methods (BADGE 10.10 h, Core-set 10.81 h) are faster than any Deep Ensemble configuration, while the most expensive (Epistemic Top-$k$ 14.71 h, Noise-Aware 14.42 h, NN-Cov 14.31 h) run about 20% longer than Deep Ensemble's slowest. Under MC Dropout, wall-clock cost is therefore governed more by the acquisition function than by the backend choice, and no single-direction comparison between backends is supportable.

**Table S10:** Aggregate compute summary across the eight paper endpoints. Each figure is a per-acquisition-method total, pooled over eight endpoints, ten seeds, and all rounds; the wall-clock and per-round columns are the minimum–maximum across the seven acquisition methods. See the definitional notes below for exactly what each unit counts.

| Backend | Trained models | Eval passes | Acquisition passes | Wall h range | Mean s/round |
|---|---|---|---|---|---|
| MC Dropout | 877 | 112,256 | 84,192‡ | 10.10–14.71 | 41.4–60.3 |
| Deep Ensemble | 4,385 | 4,385 | 3,985 / 7,970§ | 11.19–12.22 | 45.9–50.1 |

**Definition of a trained model.** One *trained model* is one model fitted in one training event, where a training event is a single (endpoint, seed, method, round) cell. Each active-learning trajectory contains 11 training events: the initial fit on the 200-molecule labeled set plus one refit after each of the 10 acquisition rounds. Eight endpoints $\times$ 10 seeds $\times$ 11 events gives 880 nominal events per acquisition method, less the three missing MBPB round-10 cells (seeds 0, 5, and 6), i.e. 877. MC Dropout trains one model per event, so its total is 877; Deep Ensemble trains $M = 5$ members per event, so its total is $877 \times 5 = 4{,}385$. The 5$\times$ ratio is therefore exact by construction, not an empirical measurement.
**Evaluation passes.** MC Dropout requires $T_{\text{test}} = 128$ stochastic forward passes per trained model, giving $877 \times 128 = 112{,}256$. Deep Ensemble requires one deterministic pass per member, so its evaluation-pass count coincides with its trained-model count at 4,385.
**Acquisition passes, and why only Deep Ensemble spans two values.** Acquisition occurs in 10 of the 11 events per trajectory (the final round is evaluated but never acquires from), giving 797 acquisition events per method. Deep Ensemble scores the pool with its five members once per acquisition event, i.e. $797 \times 5 = 3{,}985$ passes, for Epistemic Top-$k$, Noise-Aware, COVDROP, and NN-Cov. BADGE and Core-set require a second full pass per event to extract last-layer gradient embeddings and GNN embeddings respectively, so they cost $797 \times 5 \times 2 = 7{,}970$. The Deep Ensemble entry is thus two discrete values, not a continuous range: 3,985 for four methods and 7,970 for two. All six non-random MC Dropout methods instead share one $T_{\text{acq}} = 96$ shortlist-scoring budget and are recorded identically, which is why that entry is a single point value.
‡The MC Dropout acquisition figure is an upper bound. MC Dropout run metadata does not record a per-round acquisition-pass count, so the compute report applies the $T_{\text{acq}} = 96$ budget to every recorded event, including the final evaluation-only event. On the same 797-event basis used for Deep Ensemble the figure is $797 \times 96 = 76{,}512$ shortlist passes, plus $797 \times T_{\text{screen}} = 6{,}376$ cheap full-pool screening passes. Note also that the 96 refined passes score only the shortlist of $10 \times k = 480$ candidates, whereas each Deep Ensemble pass scores the full pool, so pass counts are not directly comparable in floating-point cost between backends.
§3,985 for Epistemic Top-$k$, Noise-Aware, COVDROP, and NN-Cov; 7,970 for BADGE and Core-set.

### Validation Index

Both backend roots passed schema validation and the joint backend validator. Shared split artifacts, paired bootstrap outputs, actionability diagnostics, compute summaries, and learning-curve data were regenerated from the validated paper-ready outputs. The full per-seed split audit, prediction/acquisition schemas, paired confidence intervals, learning-curve summaries, endpoint-by-method actionability diagnostics, and endpoint-by-method compute records are available in the accompanying analysis output directory. This separation keeps the print supplement within the requested length while preserving reproducibility.
